# Impairments in Hippocampal Place Cell Sequences during Errors in Spatial Memory

**DOI:** 10.1101/2020.04.20.051755

**Authors:** Chenguang Zheng, Ernie Hwaun, Laura Lee Colgin

## Abstract

Theta and gamma rhythms temporally coordinate sequences of hippocampal place cell ensembles during active behaviors, while sharp wave-ripples coordinate place cell sequences during rest. We used a delayed match-to-place memory task to investigate whether such coordination of hippocampal place cell sequences is disrupted when memory errors occur. As rats approached a learned reward location, place cell sequences represented paths extending toward the reward location during correct trials. During error trials, paths coded by place cell sequences were significantly shorter as rats approached incorrect stop locations, with place cell sequences starting at a significantly delayed phase of the theta cycle. During rest, place cell sequences replayed representations of paths that were highly likely to end at the correct reward location during correct but not error trials. The relationship between place cell sequences and gamma rhythms, however, did not differ between correct and error trials. These results suggest that coordination of place cell sequences by theta rhythms and sharp wave-ripples is important for successful spatial memory.

## INTRODUCTION

An important question in systems neuroscience is which hippocampal network patterns are critical for successful spatial memory. There are three major patterns in hippocampal local field potentials (LFPs) in freely behaving rodents: theta rhythms, sharp wave-ripples (SWRs), and gamma rhythms. Each of these patterns has been hypothesized to play an important role in learning and memory^1^. Yet the extent to which these network dynamics go awry at times when memory fails is not fully understood.

Numerous studies over the past several decades have suggested that hippocampal theta rhythms are important for learning and memory^2^. One mnemonic function that has been suggested for theta is to organize ensemble activity of hippocampal neurons into a compressed representation of ongoing spatiotemporal experiences (“theta sequences”)^3–10^. Theta sequences have been shown to develop with experience^7^, consistent with the hypothesis that they play a role in learning and memory. In addition, disruption of theta sequences by medial septum inactivation has been associated with impaired performance of a memory task^10^. However, it remains unknown whether theta sequences are impaired during memory errors that occur when theta rhythms are intact. One important study reported that theta sequences at the start of an animal’s journey predict a distance that extends further ahead of the animal when they are traveling to a goal that is further away^8^. However, this study reported no difference in the distance of theta sequences’ predicted paths as animals approached a goal destination. Moreover, this study showed that theta sequences could reflect rats’ future choices but did not address the question of whether paths coded by theta sequences are related to performance on memory tasks.

Previous studies have also investigated sequences of place cell firing that occur during SWRs in sleep and awake rest^11–14^. Such sequences “replay” representations of trajectories from earlier active behaviors on a faster time scale and have been proposed to be important for memory consolidation, memory retrieval, and planning of future trajectories^13, 14^. In line with this idea, blockade of neuronal activity during SWRs has been shown to impair memory performance^15–17^. Another study reported that, during periods of immobility in a spatial memory task, place cell ensembles tended to replay trajectories that ended at learned goal locations^18^. A recent study reported that replay of forward-ordered sequences of locations representing animals’ future paths toward goal locations was impaired on trials when animals made errors^19^. However, this replay bias was observed during rest periods at a reward well in a spatial working memory task. It is unclear whether similar results would be observed during rest periods prior to memory retrieval errors on tasks involving a significant delay (i.e., tens of seconds) between training and testing. This is an interesting question that deserves attention because such tasks have been shown to require an intact hippocampus^20, 21^.

A number of rodent studies have also related gamma rhythms to performance on mnemonic tasks^22–30^. In addition, recent studies have shown that low (∼25-55 Hz) and high (∼60-100 Hz) frequencies of hippocampal gamma activity are associated with CA3 and entorhinal cortex inputs to subfield CA1^31–34^, respectively, raising the possibility that “slow” and “fast” gamma subtypes facilitate transmission of different inputs to CA1 place cells. Thus, it is possible that slow and fast gamma coordination of place cell spike sequences would be abnormal during unsuccessful spatial memory operations.

In this study, we recorded from hippocampal subfield CA1 of rats performing a delayed match-to-sample task to assess whether spatial trajectories represented by place cell sequences differed between correct and incorrect trials. We hypothesized that incorrect trials would be associated with deficient coordination of place cell sequences by theta, gamma, and sharp wave-ripples, reflecting error modes of the hippocampal memory network and disrupted access to memory storage. To test this hypothesis, we recorded hippocampal place cell ensembles and LFP patterns as animals actively performed the delayed match-to-sample task and during rest periods. As rats learned reward locations, hippocampal place cell sequences developed that represented paths extending ahead of their current locations. Moreover, as rats approached the end of their trajectory toward their stop location, place cell sequences represented longer trajectories on correct trials than on error trials. During rest periods, place cell ensembles showed high fidelity replay of task trajectories during both correct and error trials. However, place cell sequences exhibited a significant bias to replay a path that terminated at the correct reward location only during correct trials. For both slow and fast subtypes of gamma, no differences in gamma phase coordination of spikes were observed between correct and error trials. Taken together, these results suggest that correct behavioral performance on a spatial memory task is associated with the development of coordinated sequences of place cells that represent trajectories extending toward a learned reward location.

## RESULTS

### Behavioral task and performance

To investigate whether network dynamics in the hippocampus are abnormal when memory fails, we used a circular track version of a delayed match-to-sample task (Fig. 1a). Each day, recordings began with 4-6 “pre-running” trials. In pre-running trials, rats simply completed laps around the track without receiving any rewards. This stage of the task had no memory component but allowed time for experience-dependent changes in hippocampal rhythms^35, 36^ to reach a stable level and provided a data set that was used to cross-validate a Bayesian decoder (Supplementary Fig. 1). The pre-running trials were also used to verify that rats did not prefer to spend time at particular locations such as the reward location from the previous day. In the next stage of the task, rats performed 8 trials consisting of sample-test pairs. In the sample-test trials, a reward location on the track was pseudo-randomly selected each day and remained constant across trials so that learning across trials could be assessed. During the sample phase, the reward location was marked with a visual cue, and the rat was rewarded for stopping at the marked location. In the test phase, the cues were removed such that a rat was required to recall the correct reward location from memory and stop there in order to receive a reward. Sample and test phases were separated by a 30 second delay, and inter-trial intervals were approximately 1 minute. Five minutes after sample-test trials were completed, 4-6 “post-test” trials were performed. Post-test trials were identical to the test phase in the previous stage (i.e., animals were rewarded for stopping at the same unmarked reward location). Post-test trials were included to verify that rats stored a memory of the day’s reward location. Behavioral performance data from the task showed that rats (n = 4) learned the correct reward location within the first few trials and continued to perform significantly above chance during post-test trials (Fig. 1b). Across all recording sessions and rats, the most common errors were stopping one location ahead or behind the correct reward location (Fig. 1c).

**Fig. 1.**
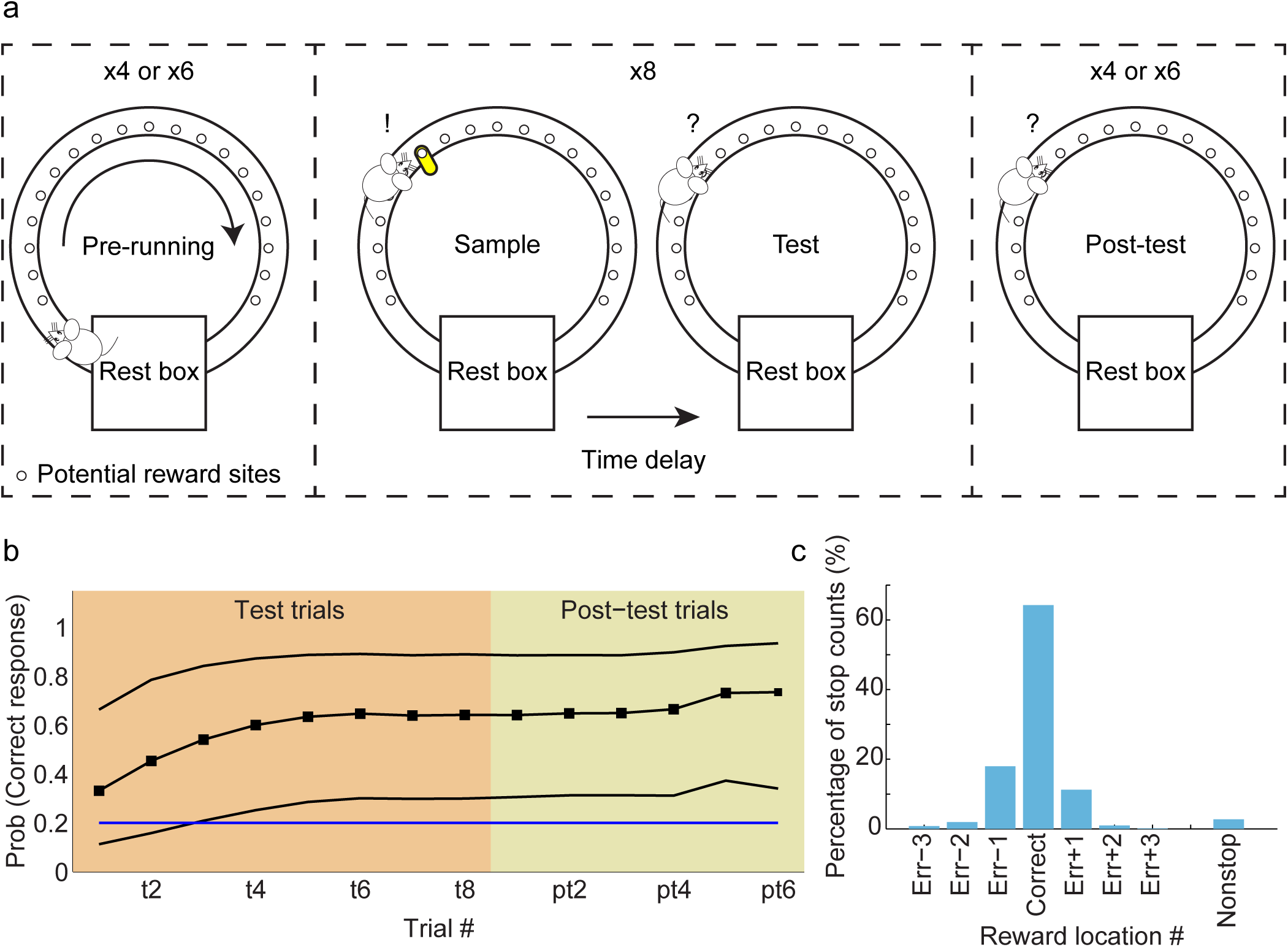
Behavioral task and performance. **a**, Schematic of delayed match-to-sample task design. Each session consisted of pre-running trials, sample-test trials, and post-test trials. In pre-running trials, rats ran 4 or 6 laps unidirectionally on a circular track without receiving any reward. One of the potential reward sites was pseudo-randomly chosen and marked with a visual cue in sample trials. The marker was removed in test and post-test trials. Rats received a reward if they stopped at the goal location. The goal location remained constant within a session. **b**, Behavioral performance across test and post-test trials is shown. Black solid lines represent the mean probability of making a correct choice across recording sessions, bounded by 95% confidence intervals. The blue horizontal line indicates chance performance. **c**, Proportion of trials in which rats stopped at different locations in test and post-test trials pooled across all recording sessions.

Importantly, no differences were observed in running speeds between correct and error trials (repeated measures ANOVA, main effect of trial types: F(1,42) = 1.2, p = 0.3). Moreover, rats were not found to linger for a longer amount of time in the rest box before beginning to run on the test phase of error trials (paired t-test: T(42) = 0.6, p = 0.5), nor did they remain at the reward location for a longer amount of time during the sample phase of correct trials (paired t-test: T(42) = 1.5, p = 0.1). During the pre-running phase of the task, there was no difference in rats’ preferences to spend time at the reward location from the previous day, the current day’s reward location, or the current day’s erroneous stop locations (repeated measures ANOVA, F(2,84) = 0.4, p = 0.6). These results suggest that errors are not explained by differences in rats’ behaviors or biases for particular locations and may instead involve impaired network dynamics in the hippocampus.

### Place cell sequences representing paths to the reward location developed with experience

A previous study showed that theta-coordinated sequences of place cells emerge with experience on a novel linear track, a task that does not require memory^7^. Thus, we hypothesized that sequences of place cells representing paths toward a reward location in a spatial memory task would develop as rats learned the reward location across trials. To test this hypothesis, we recorded ensembles of place cells in hippocampal subregion CA1 of rats performing the above-described delayed match-to-sample task. We then used a Bayesian decoding approach (see Methods; Supplementary Fig. 1) to identify trajectories represented as rats approached the reward location in sample and test phases of sample-test trials. We found that place cell sequences’ representations of paths extending ahead of the animal increased across trials for both sample and test phases (Fig. 2a; repeated measures ANOVA, main effect of trial number: F(7,231) = 7.7, p = 2 x 10^-8^). In contrast, place cell ensemble representations of locations behind the animal remained constant across trials (Fig. 2b; repeated measures ANOVA, significant interaction between location type (i.e., ahead and behind) and trial number: F(7,497) = 5.5, p = 4 x 10^-6^; no main effect of trial number for behind locations: F(7,266) = 1.1, p = 0.4). These results support the conclusion that coordinated sequences of place cells that predict paths toward upcoming reward locations during theta-related behaviors develop as animals learn reward locations across trials.

**Fig. 2.**
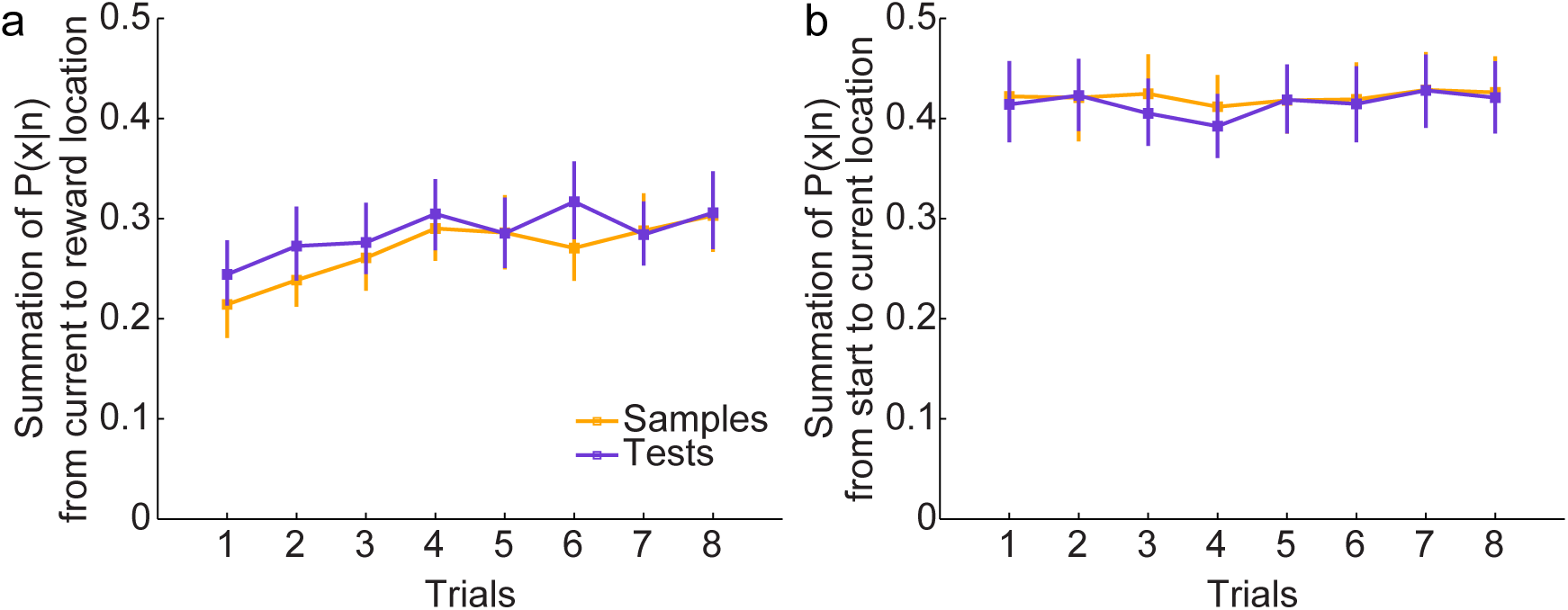
Predictive firing in place cell ensembles developed with learning. A Bayesian decoder (see Methods) was applied to ensemble spiking activity to estimate posterior probability distributions across sample-test trials. **a**, The sum of posterior probability (P(x|n)) extending ahead of rats (i.e., from a rat’s current position to the reward location) as rats ran toward their stop location is shown for each trial. The sum significantly increased across trials (repeated measures ANOVA, main effect of trial number: F(7,231) = 7.7, p = 2 x 10^-8^). **b**, The sum of posterior probability behind rats’ current positions is shown across sample-test trials. The posterior probability behind rats’ current positions (i.e., extending from the beginning of the track to a rat’s current position) was summed for each lap. In contrast to the results reported in **a**, the sum of posterior probability behind rats’ current positions did not significantly change across trials (repeated measures ANOVA, significant interaction between location type (i.e., ahead and behind) and trial number: F(7,497) = 5.5, p = 4 x 10^-6^; no main effect of trial number for behind locations: F(7,266) = 1.1, p = 0.4). Posterior probability values appear higher for behind positions than for ahead positions; note that this does not mean that theta sequences are longer in the behind direction. Rather, it means that the probability that the animal is in those locations based on the ensemble spiking patterns is higher. Posterior probability values for decoded sequences would be expected to be higher for current locations and just visited locations than for locations extending ahead of the animal because place cells fire at higher rates at the end of their place fields than in the early, negatively skewed portion of their place fields^49^. The posterior probability is calculated based on the probability for spikes to occur at a given location (i.e., P(n|x), which is determined from the firing rate maps). Error bars indicate 95% bootstrapped confidence intervals.

### Place cell sequences represented shorter trajectories as rats approached an incorrect stop location during error trials

It has been hypothesized that organized sequences of place cells firing within theta cycles are important for retrieving previously stored representations of upcoming trajectories^4, 6, 37^ or encoding of current trajectories^3, 4, 6, 38^. Therefore, we hypothesized that sequences of place cells activated during theta-related behaviors would differ based on memory performance in a delayed match-to-sample task. To quantify represented trajectories during correct and error trials, we decoded place cell ensemble activity as rats ran toward their eventual stop locations (i.e., the stop location was the same as the correct reward location on correct trials but corresponded to animals’ incorrect stop locations on error trials). We then detected epochs that showed significant sequential structure and measured these sequences’ slopes as an estimate of distance and direction of decoded paths (see Methods). We hypothesized that coordinated place cell sequences predicting paths to the upcoming stop location would be more prominent during the test phase of correct trials, considering that place cell sequences developed a representation of upcoming paths as the reward location was learned across trials. In line with this hypothesis, we found that slopes of sequences detected as rats approached their stop locations were greater for correct test trials than for error test trials (Fig. 3, Supplementary Fig. 2; ANOVA, main effect of trial type (i.e., correct vs. error): F(1,2826) = 15.0, p = 1.1 x 10^-4^).

**Fig. 3.**
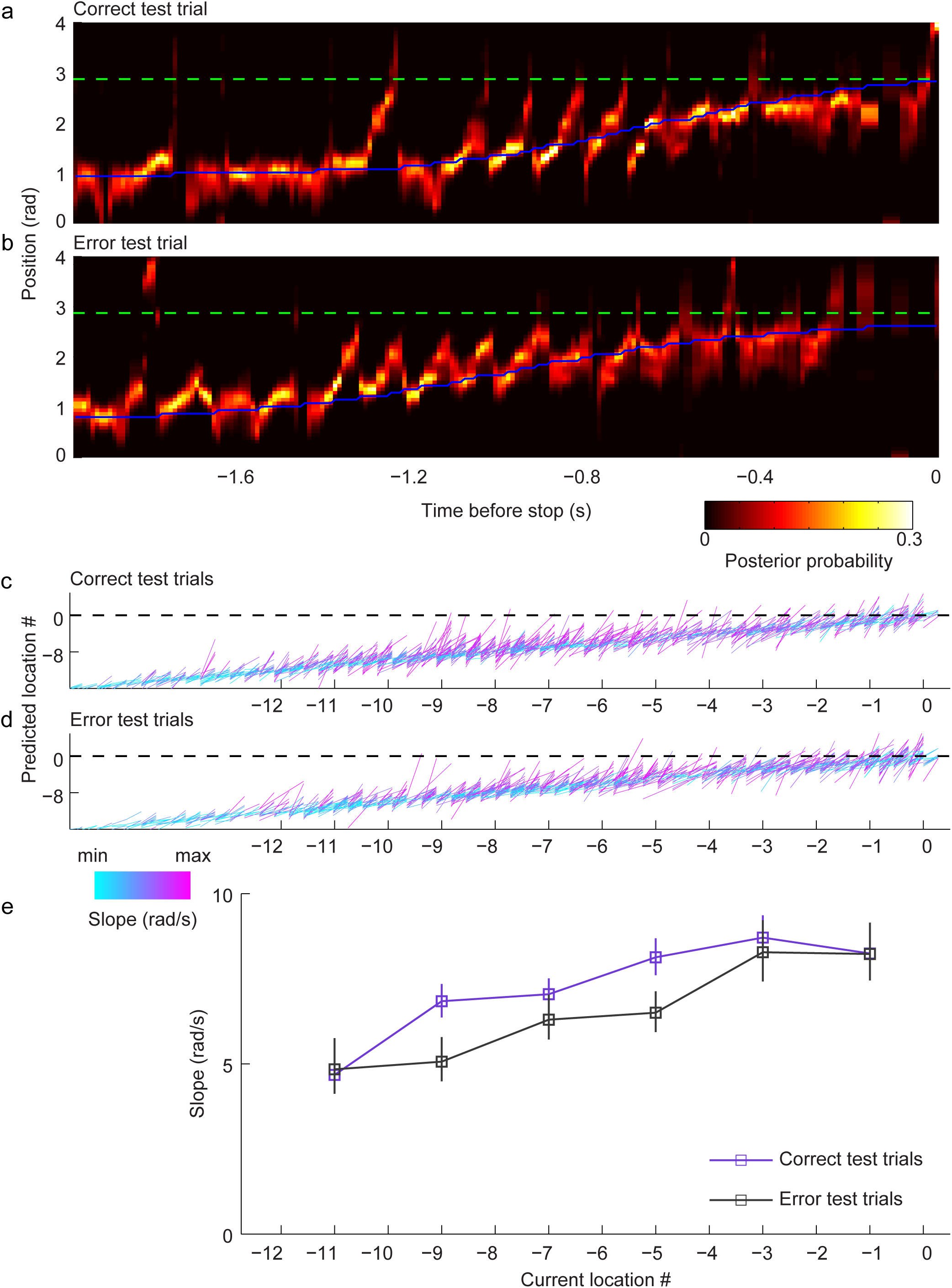
Place cell sequences exhibited steeper slopes in correct test trials than in error test trials. **a**, An example color-coded posterior probability distribution across time from an example test trial in which the rat stopped at the correct reward location. The green dashed line marks the correct goal location, and the blue solid line represents the actual position of the rat at each time point. Note that the posterior probability sweeps ahead of the rat’s current position toward the reward location as the rat approached the reward location. Trajectories were aligned according to the time when the animal stopped (time = 0). **b**, Same as **a** but from an example error test trial in the same recording session. The rat stopped one location (or ∼0.2 rad) earlier than the correct goal location in this trial (see Supplementary Figure 2d for an example error trial in which the rat stopped one location too late). Note that the posterior probability in this trial did not extend as far ahead of the rat’s current position as the posterior probability in **a**. **c-d**, For purposes of illustration and comparison, place cell sequences in correct test trials (**c**) were randomly down-sampled to match the number of place cell sequences in error test trials (**d**). Fitted lines of detected place cell sequences in correct (**c**) and error (**d**) test trials are shown across location numbers as rats approached their stop location (indicated as location 0; same location as correct goal location for correct trials). The x and y coordinates of each sequence’s fitted line indicate current position and predicted position (i.e., maximum posterior probability), respectively. The slope of the fitted line for each sequence is shown color-coded. The black dashed line signifies the rats’ stop location. Decoded locations within a sequence that extended beyond the stop location were included as part of that sequence, and locations in the sequence that extended beyond the stop location were included in the slope calculation. **e**, Mean slopes of the lines that were fit to detected place cell sequences are shown across current locations as rats approached their stop location. Slope measurements are shown plotted for the center of each location bin (i.e., locations −12 to −10 plotted at location −11, locations −10 to −8 plotted at location −9, etc.). Error bars indicate 95% bootstrapped confidence intervals.

Sample phases in trials after the initial trial likely involve not only encoding of the marked reward location but also recall of the rewarded location from previous trials. Therefore, we also analyzed slopes of sequences detected during the sample phase of the task to determine whether sequences from correct trials were different than sequences during error trials. Slopes of sequences as animals approached stop locations during the sample phase were also different between correct and error trials, with greater slopes again observed during correct trials (Fig. 4; main effect of trial type: F(1,2838) = 15.9, p = 6.9 x 10^-5^).

**Fig. 4.**
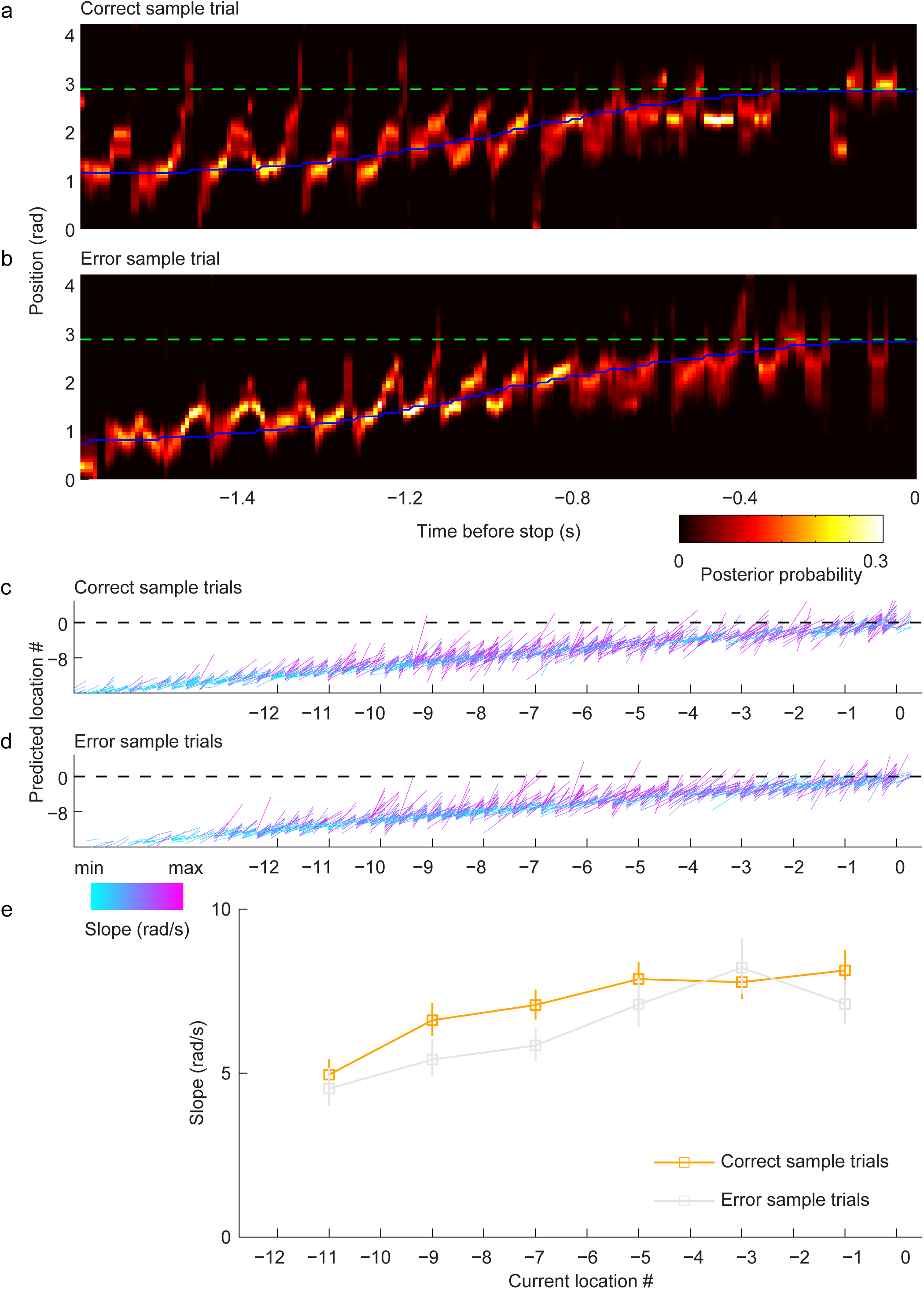
Place cell sequences exhibited steeper slopes in correct sample trials than in error sample trials. **a-e**, Same as **a-e** in Fig. 3 but for sample trials. Note that correct and error trials were defined by rats’ responses in subsequent test trials.

Slope provides a measure of spatial shifts in decoded sequences (i.e., length of represented path) within a particular amount of time (i.e., the duration of a sequence), so differences in slopes could reflect differences in the amount of spatial shift, the duration of a sequence, or both. Therefore, we also assessed whether the length of sequences’ represented trajectories and their temporal duration differed between correct and error trials. The length of represented trajectories differed significantly between correct and error trials (0.612 ± 0.414 and 0.536 ± 0.381 radians for correct and error trials, respectively; F(1,5664) = 54, p = 3 x 10^-13^), but the temporal duration of detected sequences did not (0.108 ± 0.023 s and 0.109 ± 0.024 s for correct and error trials, respectively; F(1,5664) = 2.2, p = 0.1). These results imply that correct recall of trajectories toward learned reward locations requires place cell sequences that code a relatively long trajectory extending ahead of the animal toward the correct stop location. These results were not explained by differences in running speeds during the approach to the stop location because running speeds during the approach to the stop location did not differ between correct and error trials (Supplementary Fig. 3).

The above-described results raise the question of what mechanisms underlie the longer trajectories represented by place cell sequences as rats approached their stop location during correct trials compared to error trials. To gain some potential insight into this question, we looked at the slopes of all sequences (Fig. 3c-d and Fig. 4c-d) and at posterior probability distributions for individual trials (examples shown in Fig. 3a-b, Fig. 4a-b, and Supplementary Fig. 2). Sequences appeared to begin slightly earlier as rats approached their stop location during correct trials, providing a possible explanation for the correct trials’ longer decoded trajectories. To test this possibility, we assessed the theta phases at which place cell sequences’ spikes occurred as rats approached their stop locations during correct and error trials (Fig. 5). Theta phase coding of position was apparent across all trial types. That is, place cells that had place fields at relatively early and relatively late locations fired at relatively early and relatively late theta phases, respectively (see “Categorization of place cell locations” subsection of Methods). Yet, the theta phase distributions differed significantly in an interesting way (non-parametric circular statistical test for equal medians across the 4 trial types: p = 0.004). Place cells that represented relatively early locations in the trajectory were more likely to fire at an earlier phase of the theta cycle during correct trials than during error trials (non-parametric circular statistical test for equal medians for correct and error trials, p = 0.001). Results were similar for both sample and test phases (correct vs. error for sample phase only: p = 0.004; correct vs. error for test phase only: p = 0.05). Similar results were not obtained for place cells with place fields at relatively late locations along the trajectory (non-parametric circular statistical test for equal medians across the 4 trial types: p = 0.4; correct vs. error trials: p = 0.2; correct sample vs. error sample: p = 0.3; correct test vs. error test: p = 0.3). These results suggest that place cell sequences code longer trajectories extending toward the stop location on correct trials because place cells that start the sequence begin firing at a significantly earlier theta phase during correct trials than during error trials. That is, place cells that coded locations early in a rat’s trajectory were more likely to start firing at an earlier phase of the theta cycle on correct trials than on error trials.

**Fig. 5.**
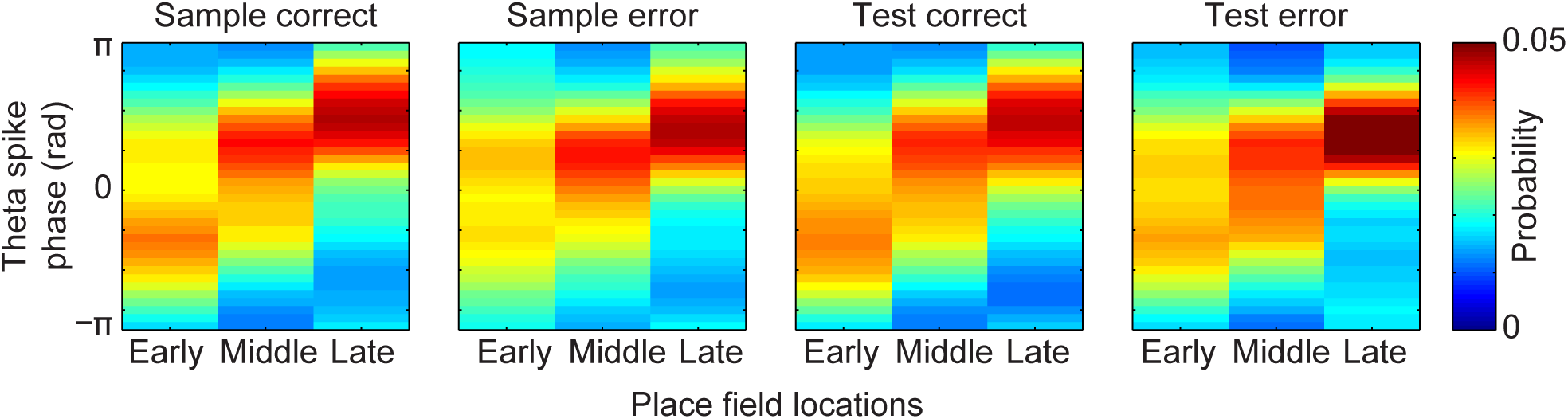
Place cells active at the beginning of the trajectory toward the reward fired at earlier theta phases in correct trials than in error trials. The probability of observing a spike at different theta phases for spikes from place cells with fields located at early, middle, or late locations in the trajectory toward the reward is shown for each trial type. Phase estimates were only obtained from spikes within the place cell sequences detected from 10 to 5 locations before the goal.

We preferred to define sequences based solely on ensemble spiking (i.e., epochs in which spikes from place cell ensembles showed significant sequential structure), regardless of the properties of hippocampal rhythms present during those epochs. This allowed us to perform sequence analyses without making assumptions about the theta phase at which sequences began and ended or about the maximum duration of sequences. Nevertheless, we also performed analogous analyses on sequences that were detected within individual theta cycles. Sequences detected within individual theta cycles were found to exhibit greater slopes in correct trials than in error trials as animals approached stop locations during the test phase of the task (main effect of trial type (i.e., correct vs. error): F(1,1101) = 65.1, p = 0.003). Slopes of sequences detected within individual theta cycles were not significantly different between correct and error trials as animals approached their stop locations during the sample phase (main effect of trial type: F(1,1092) = 1.1, p = 0.3). However, this lack of significance may have been due to reduced statistical power because the number of sequences decreased when sequences were detected within individual theta cycles. In support of this assumption, we found that slopes of sequences detected within individual theta cycles were significantly different between correct and error trials when sequences from the sample and test phases of the task were combined (main effect of trial type: F(1, 2193) = 7.9, p = 0.005; no significant interaction between trial type (i.e., correct vs. error) and task phase (i.e., sample vs. test): F(1,2193) = 1.6, p = 0.2).

For the results described above, only sequences that represented locations extending ahead of the animal (i.e., sequences with positive slopes) were included. Similar results were obtained when sequences with positive slopes and sequences with negative slopes (i.e., sequences that do not represent locations extending ahead of the animal) were combined together (main effect of trial type (i.e., correct vs. error): F(1,4384) = 28.3, p = 1.1 x 10^-7^ for test phase and F(1,4449) = 24.5, p = 7.7 x 10^-7^ for sample phase). Significant differences between correct and error trials were also observed when sequences with negative slopes were analyzed separately (Supplementary Fig. 4; ANOVA, main effects of trial type: F(1,1599) = 7.3, p = 0.007 for sample phase; F(1,1546) = 13.2, p = 2.9 x 10^-4^ for test phase; F(1,3145) = 20.0, p = 8.1 x 10^-6^ for sample and test phases combined). The proportion of positive slope to negative slope sequences was significantly higher when sequences were detected based solely on sequential structure in ensemble spike trains than when sequences were detected within individual theta cycles (5688 and 3169 positive and negative slope sequences, respectively, detected from continuous ensemble spiking; 2565 and 2250 positive and negative slope sequences, respectively, detected within individual theta cycles; χ^2^(1) = 156.3, p = 7.3 x 10^-36^).

### Replay during awake rest develops a bias toward the reward location during correct but not error trials

The above results suggest that organized sequences of place cell activity during theta-related behaviors may be important for learning and correct performance of memory tasks. Replay during SWRs is also thought to play a key role in learning and memory, and blocking neuronal activity during SWRs has been shown to decrease performance on memory tasks^15–17^. Thus, we first hypothesized that the fidelity of replay sequences would be poorer during error trials compared to correct trials. To assess replay of trajectories, we employed a Bayesian approach to decode place cell ensemble spiking activity recorded when rats were resting (see Methods). In contrast to our original hypothesis, we found that place cells replayed trajectories on the circular track with high fidelity during the rest periods of both correct and error trials (Fig. 6) for both forward and reverse replay events (Supplementary Fig. 5).

**Fig. 6.**
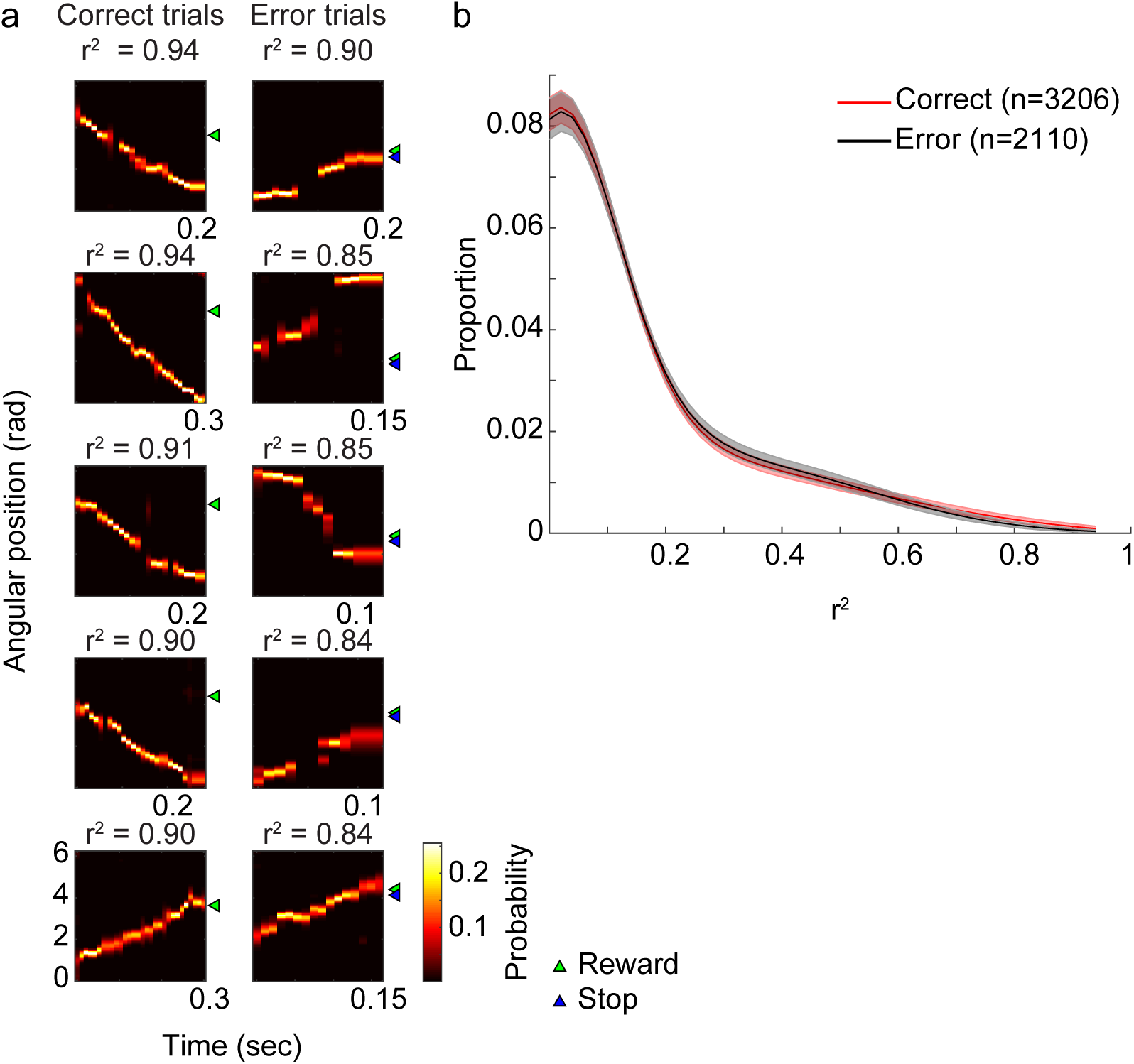
Replay quality in the rest box was similar between correct and error trials. **a**, The SWR events with the top 5 r^2^ values in each trial type are shown for correct and error trials. The color scale indicates posterior probability from a Bayesian decoder (see Methods). SWR events were detected while rats were in the rest box between sample and test trials. The green and blue triangles mark reward and stop positions, respectively, from each trial. **b**, Distributions of r^2^ values for all detected SWR events in correct and error trials. Shaded error bars indicate 95% bootstrapped confidence intervals.

A previous study reported that replay of forward paths toward a goal location in a working memory task was reduced in error trials compared to correct trials^19^. Here, we hypothesized that trajectories represented during replay also differ between correct and error trials in a delayed match-to-sample task. We examined the content of replayed trajectories during rest periods of sample/test trials and found differences between correct and error trials. Specifically, a bias in replay to terminate at the reward location developed across trials on correct but not error trials (Fig. 7, Supplementary Fig. 6). This bias was observed when rest periods before and after the test phase of trials were grouped together (Fig. 7) and when rest periods before and after the test phase were analyzed separately (Supplementary Fig. 6). A similar bias toward the correct reward location persisted during rest periods in correct post-test trials (Supplementary Fig. 7a-b). In contrast, there was no bias in replay observed during the pre-run period prior to learning (Supplementary Fig. 7c-d). No significant bias for replay events to start from the goal location was found (Supplementary Fig. 8). Also, there was no relationship between the representation of a particular end location during replay events in error trials and animals’ subsequent stop locations (Supplementary Fig. 9).

**Fig. 7.**
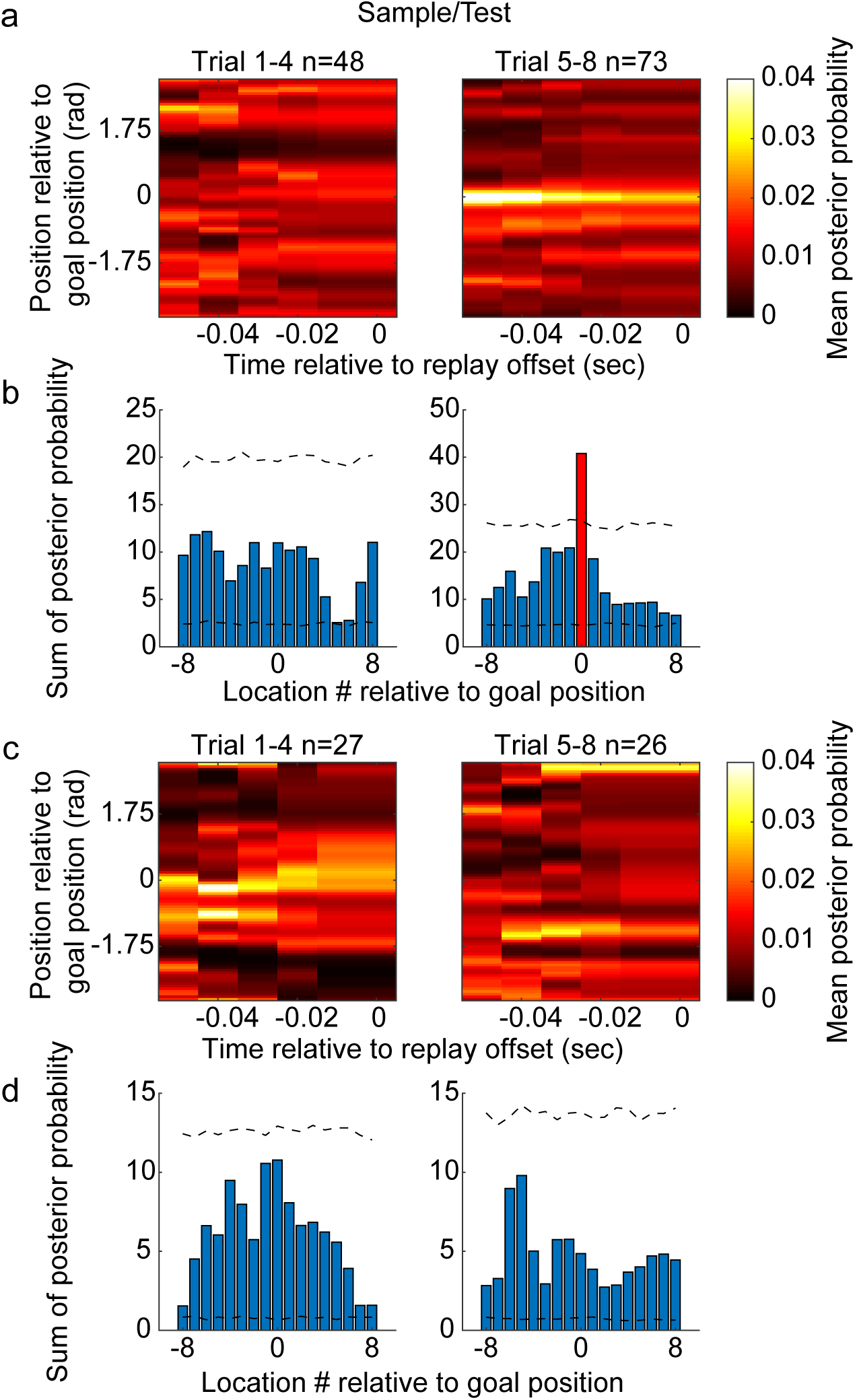
A bias for replay events to terminate at the correct goal location emerged after learning during correct trials. **a**, Mean posterior probabilities of replay events in correct trials. Replay events were aligned to the correct goal location on the vertical axis (i.e., location 0) and the time of replay event offset on the horizontal axis. **b**, The sum of posterior probability across time for each location number relative to the goal. Dashed black lines mark 95% confidence intervals of a null distribution generated by randomly shifting positions of each replay event (see Methods). A red bar marks the location number with a posterior probability sum that exceeded the corresponding 95% confidence interval (i.e., location 0, which corresponds to the correct goal location). **c-d**, Same as **a-b** but for error trials. Note that no positions during error trials showed a posterior probability sum that was greater than the corresponding confidence interval.

### Errors were not associated with deficits in gamma phase modulation of place cell spikes

Besides theta rhythms and SWRs, gamma rhythms have also been hypothesized to play a role in learning and memory^22–30^. Previous studies have also suggested that “slow” (∼25-55 Hz) and “fast” (∼60-100 Hz) gamma rhythms are associated with different inputs to the hippocampus^31–34^. Specifically, detection of fast and slow gamma activity in CA1 has been associated with inputs from medial entorhinal cortex and CA3, respectively. In our earlier work, we showed that spikes at successive slow gamma phases tended to code consecutive spatial positions in a sequence of locations^9^. Based on these results, we hypothesized that slow gamma phase coding provides a mechanism for predicting sequences of upcoming locations and that this mechanism is important for spatial memory recall. To test this hypothesis, we compared distributions of slow gamma phases of spike times as rats approached their stop locations during correct and error trials (Figure 8a, c; Supplementary Fig 10a). Shifts in slow gamma phases of place cell spike times across cycles appeared virtually identical across correct and error trials for both sample and test phases of the task (Fig. 8a, c). Moreover, as rats approached their stop location, robust differences in slow gamma power were not apparent between correct and error trials (Supplementary Fig. 11). These findings are inconsistent with our original hypothesis that a failure of slow gamma phase coding of sequential positions underlies errors in spatial memory recall. In our earlier report^9^, we also hypothesized that fast gamma phase-locking of spikes promotes real-time memory encoding. If so, then fast gamma phase-locking of spikes should be impaired during error trials. To investigate this possibility, we compared distributions of fast gamma phases of spikes as rats approached their stop locations during correct and error trials Fig. 8b, d; Supplementary Fig. 10b). Place cell spikes were similarly phase-locked to fast gamma during the sample and test phases of correct and error trials (Fig. 8b, d). Moreover, no significant differences in fast gamma power between correct and error trials were observed as rats approached their stop location (Supplementary Fig. 11). Taken together with the results reported above, these findings suggest that errors in a spatial memory task are associated with deficient place cell sequences but that deficits are not associated with abnormal coordination by slow or fast gamma rhythms.

**Fig. 8.**
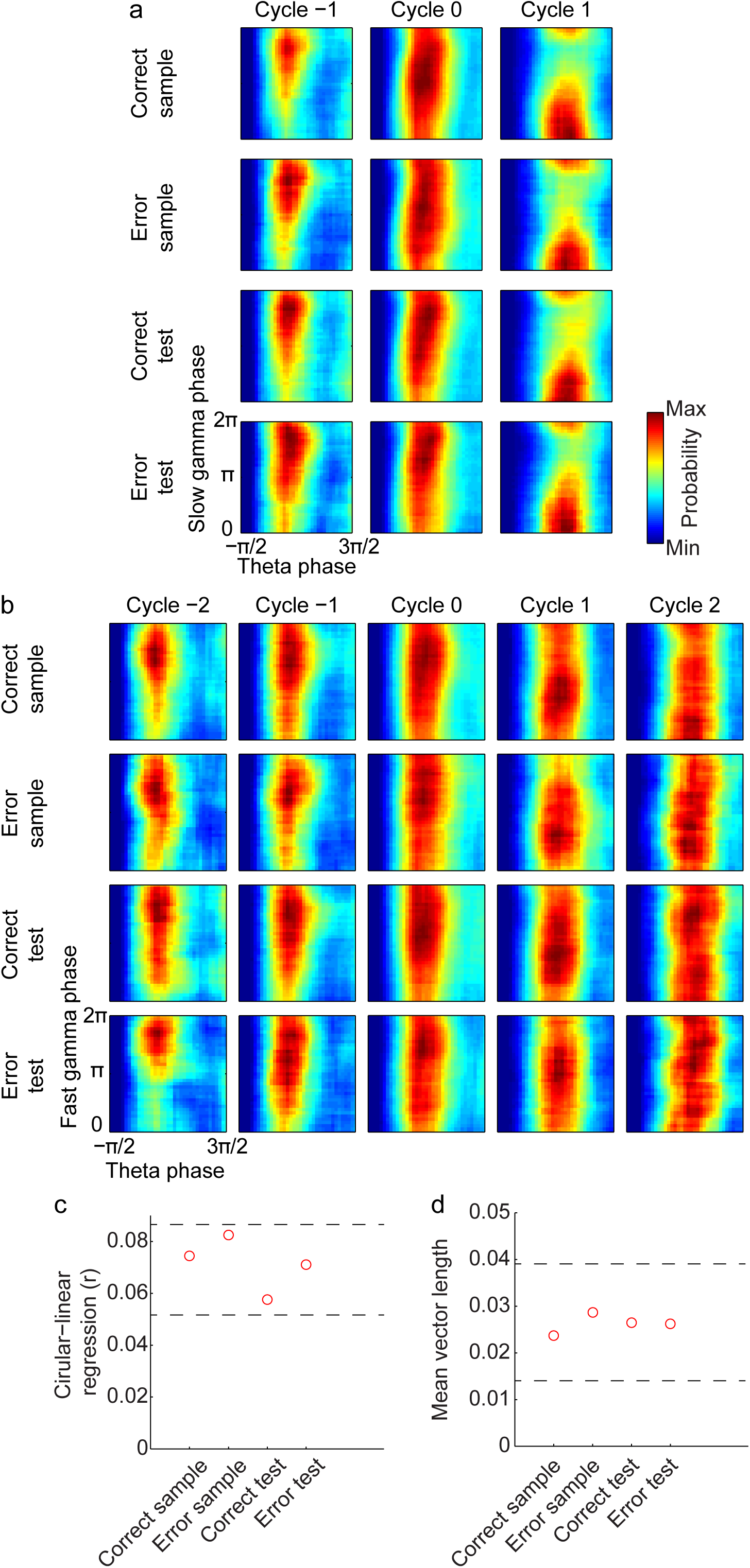
Gamma phase modulation of place cell spikes was similar between different trial types. **a**, Probability distributions of slow gamma phases of spikes across slow gamma cycles within place cell sequences for each trial type. Slow gamma phases of spikes shifted systematically across successive slow gamma cycles in all trial types. **b**, Probability distributions of fast gamma phases of spikes across successive fast gamma cycles within place cell sequences for each trial type. Fast gamma phase distributions remained similar across successive fast gamma cycles and across trial types. **c**, Circular-linear regression between slow gamma phases of spikes and cycle number for each trial type indicate that slow gamma phases of place cell spikes shifted across successive gamma cycles during both correct and error trials. **d**, Mean vector lengths of fast gamma phases of place cell spikes for each trial type, pooled across all fast gamma cycles. Note that mean vector lengths of fast gamma phase distributions were similar across trial types. Black dashed lines in **c-d** mark 95% confidence intervals of a null distribution generated by shuffling trial types.

## DISCUSSION

In a delayed match-to-sample task in which a different reward location was learned each day, organized sequences of place cells represented extended paths as rats moved toward the reward location on correct trials. In contrast, place cell sequences represented shorter paths as rats approached incorrect stop locations on error trials. Sequence impairments were also observed during rest periods of error trials. Specifically, as rats rested between runs, place cell sequences replayed trajectories that terminated at the learned reward location during correct trials but not during error trials. Contrary to a hypothesis proposed in our earlier study^9^, gamma phase coordination of place cell sequences during error trials did not differ from phase coordination during correct trials. These results provide novel insights about how hippocampal network dynamics differ between correct and incorrect performance of a spatial memory task.

### Memory errors and aberrant place cell sequences during theta-related behaviors

Prior studies have shown that theta sequences are not initially present in a novel environment but develop as an environment becomes familiar^7^ and that place cells’ order in theta sequences reverses when animals run backward^39^. These prior results suggest that organized sequences of place cells emerge with experience and reflect changes in behaviors. However, these studies did not address the question of whether place cell ensembles’ representations of paths extending ahead of the animal during the approach to a goal destination are related to correct memory recall. To address this question, we trained rats to perform a spatial delayed match-to-sample task. We found that organized sequences of place cells developed representations of paths extending toward a goal location as animals learned that location. We also found that place cell sequences represented extended trajectories toward the learned goal location as rats approached that location during correct trials and shorter trajectories as animals approached incorrect stop locations during error trials. Errors included trials in which animals stopped at locations both before and after the correct goal location. These findings imply that coordinated, experience-dependent sequences of place cells are important for correct memory recall during active performance of spatial memory tasks. It is possible that the transmission of these predictive sequences from the hippocampus to downstream structures (e.g., prefrontal cortex and nucleus accumbens) during active theta-related behaviors allows hippocampal memory operations to guide ongoing behavior. In line with this possibility, correlated activity between pairs of prefrontal neurons increases during periods when theta coherence between hippocampus and prefrontal cortex is high^40^, and spatial appetitive memory requires projections from CA1 to nucleus accumbens^41^.

A previous related study showed that place cell sequences represented trajectories that extended a greater distance ahead of rats’ current position when they began their journeys toward more distant goals^8^. However, these prior results do not explain the findings reported here. The previous findings were observed as rats started their trajectory (see their Figure 3) but not as rats approached their ultimate stop locations (see their Figure 4b and 4d). In contrast, here we show that place cell sequences represented longer paths as rats approached a learned goal location on correct trials but not as they approached an incorrect stop location on error trials (Figures 3-4). Also, the task used in the study by Wikenheiser and Redish^8^ did not permit comparison of place cell sequences from correct and error trials because rats headed toward marked feeder locations on all trials, and there was no distinction between trials based on memory performance.

The current findings suggest new hypotheses regarding the question of why errors in this task occurred. Error trials were associated with place cell sequences that coded shorter trajectories as rats approached their incorrect stop locations (Figures 3-4). Also, place cells that coded locations early in a trajectory in error trials fired at significantly later phases of the ongoing theta cycles (Figure 5). These results suggest that the hippocampus was unable to retrieve representations of extended trajectories because the place cells that represented recent and current locations (i.e., the place cells that triggered the read-out of the sequence) started firing too late. Such delayed firing may only allow part of a sequence to be read out before a strong discharge of hippocampal interneurons occurs^42^ and terminates the sequence. Delayed firing could reflect insufficient excitatory drive to the network. Insufficient excitatory drive could result from reduced neuromodulatory (e.g., cholinergic) inputs when insufficient attention is paid to the task. Or, it could perhaps result from insufficient synaptic potentiation in connections between cells coding the trajectory.

### Memory errors and place cell sequences during awake rest

A previous study involving rats performing a spatial alternation task showed that place cell co-activation during SWRs was greater during rests preceding correct trials compared to incorrect trials^43^. A subsequent study showed that place cell ensembles during rest replayed trajectories that resembled those that an animal subsequently traversed to a goal location in an open field spatial memory task^18^. A recent study further reported that forward replay of place cell sequences representing animals’ future trajectories toward a goal location in a spatial working memory task developed with learning and was impaired when animals made errors^19^. Our findings are consistent with these earlier findings but also provide new insights. We first show that place cell ensembles recorded during rest periods in a delayed match-to-sample spatial memory task developed a bias to replay representations of trajectories toward a reward site as animals learned the reward site. Furthermore, this bias was not apparent in the rest periods of trials in which animals failed to remember the correct reward site. These findings suggest that replay of place cell ensemble representations of specific trajectories to previously learned locations is necessary for successful memory recall. This hypothesis is consistent with results showing that disruption of awake SWRs as animals performed a spatial memory task impaired behavioral performance^17^. Future experiments involving real-time detection and manipulation^44^ of replay of specific trajectories will be important to test the hypothesis that replay of particular learned trajectories is required for subsequent memory retrieval.

### Relationship between place cell sequences during active behaviors and rest

The present results do not address the question of whether experience-dependent formation of place cell sequences that represent trajectories in real-time during theta-related behaviors leads to temporally compressed versions of the same sequences during SWR-associated replay. Other studies shed light on this question. Experience-dependent replay during awake rest and coordinated theta sequences have been reported to emerge at the same developmental stage (i.e., ∼P25)^45, 46^. It is likely that the coordinated sequences are initially formed during theta and subsequently activated on a faster time scale during SWRs. Support for this idea was provided by a recent study that recorded place cells ensembles in rats that were passively moved through space^47^. This passive movement served to successively activate place cells while disrupting formation of coordinated theta sequences. Importantly, passive transport and associated suppression of theta sequences blocked replay in subsequent sleep, supporting the conclusion that linking sequences of place cells during theta is required for subsequent expression of place cell sequences during SWRs. Still, theta sequences are restricted to representation of relatively short ongoing trajectories. On the other hand, SWR-associated replay can include spatially remote and extended trajectories^12, 48^. Thus, SWR-associated replay is likely necessary to link multiple sequences together in a temporally compressed manner during formation of comprehensive spatial and episodic memories.

### Slow gamma phase coding and fast gamma phase-locking remained intact during memory errors

In our earlier work, we showed that place cell spikes coding successive locations within relatively short trajectories occurred at successively increasing slow gamma phases^9^. We hypothesized that this slow gamma phase coding was an essential mechanism for retrieving representations of upcoming spatial trajectories. Our present results replicated the finding of slow gamma phase coding of locations but failed to show differences in slow gamma phase coding between successful memory and memory errors. These results are inconsistent with our original hypothesis that errors in spatial memory retrieval are related to disrupted slow gamma phase coding of locations within a learned trajectory. Instead, our results imply that memory errors in the delayed match-to-sample memory task were related to a delay in the initiation of sequences within theta cycles, as evidenced by results showing that place cells that coded early locations in the sequence fired at significantly later theta phases during error trials. Slow gamma phase coding may simply reflect read-out of previously stored sequences, regardless of whether an animal ultimately stops at a correct goal location. Related to this, it is interesting to note that predictive sequences were also observed on error trials (e.g., Fig. 4b), although sequences represented significantly shorter trajectories on error trials. Therefore, it remains possible that abnormal slow gamma phase coding could be observed in a different type of experiment involving more severe deficits in predictive sequences.

In our previous report, we also hypothesized that fast gamma phase-locking of spikes is important for successful encoding of spatial trajectories^9^. However, in the present study, no difference in fast gamma-phase locking between correct and error trials was observed. However, the present task is not well-suited to assess memory encoding as it is unclear when encoding of the trajectory to the reward location occurs. Although the reward location was changed on each recording day, the task was performed on a track and in a room that were highly familiar. Thus, it remains possible that fast gamma coordination of place cell spikes plays a role in initial memory encoding in novel situations. In support of this idea, the increase in hippocampal fast gamma power that occurs with increasing running speed, and fast gamma power overall, were maximal in a novel environment^33^. Also, CA1 fast gamma power, CA3-CA1 fast gamma phase synchrony, and fast gamma phase-locking of place cell spikes increased during encoding of novel object-place associations^27^.

### Potential Implications for Memory Disorders

The current study adds to our understanding of the importance of coordinated sequences of place cells in the hippocampus for successful learning and memory operations. The results also shed light on how hippocampal dynamics are disrupted when memory fails. It is possible that analogously abnormal hippocampal dynamics occur in brain disorders associated with learning and memory impairments. A failure to transmit learned hippocampal sequences to downstream regions that direct behavior, such as the striatum and prefrontal cortex, may contribute to spatial memory impairments in disorders like Alzheimer’s disease.

## ACKNOWLEDGMENTS

We thank A. Akinsooto, S. Dhavala, K. Kallina, and A.J. Wright for recording drive construction, histology, and other outstanding technical support; S. Brizzolara-Dove, G. Kwong, C.G. Orozco, and D. Wehle for help with rat behavioral pre-training; and all Colgin Lab members for helpful discussions. This work was supported by: the National Institutes of Health under award numbers R01 MH102450 (to L.L.C.), National Natural Science Foundation of China grants 81870847 and 31800889 (to C.Z.), and T32 MH106454 (to E.H.).

## AUTHOR CONTRIBUTIONS

C.Z. and L.L.C. designed the experiment; C.Z. and L.L.C. carried out the electrophysiological recordings and behavioral testing; C.Z., L.L.C., and E.H. designed analyses; C.Z. and E.H. wrote analysis programs and analyzed the data; L.L.C., E.H., and C.Z. wrote the paper; and L.L.C. supervised the project. All authors discussed results.

## ONLINE METHODS

### Subjects

Four male Long-Evans rats weighing from ∼400 to 600 g were used in this study. Rats were housed in custom-built acrylic cages (40cm x 40cm x 40cm) on a reverse light cycle (Light: 8pm to 8am). The cages contained enrichment materials (e.g., plastic balls, cardboard tubes, and wooden blocks). Active waking behavior recordings were performed during the dark phase of the cycle (i.e., from 8am to 8pm). Rats were pre-trained to perform the task prior to recording drive implantation surgery. Rats recovered from surgery for at least one week before behavioral training resumed. During the data collection period, rats were placed on a food-deprivation regimen that maintained them at ∼90% of their free-feeding body weight. All experiments were conducted according to the guidelines of the United States National Institutes of Health Guide for the Care and Use of Laboratory Animals under a protocol approved by the University of Texas at Austin Institutional Animal Care and Use Committee.

### Surgery and tetrode positioning

Recording drives with 20 independently movable tetrodes were surgically implanted above the right hippocampus (anterior-posterior [AP] 3.8 mm, medial-lateral [ML] 3.0 mm, dorsal-ventral [DV] 1 mm). Bone screws were placed in the skull, and the screws and the base of the drive were covered with dental cement to affix the drive to the skull. Two screws in the skull were connected to the recording drive ground. Before surgery, tetrodes were built from 17 μm polyimide-coated platinum-iridium (90/10%) wire (California Fine Wire, Grover Beach, California). The tips of tetrodes designated for single unit recording were plated with platinum to reduce single channel impedances to ∼150 to 300 kOhms. Tetrodes were gradually lowered to their target locations. Eighteen of the tetrodes were targeted to CA1 stratum pyramidale. One tetrode was designated as a reference for differential recording and remained in a quiet area of the cortex above the hippocampus throughout the experiments. This reference tetrode was moved up and down until a quiet location was found and was continuously recorded against ground to ensure that it remained quiet throughout data collection. Another tetrode was placed in the CA1 apical dendritic layer to monitor and record local field potentials (LFPs) in the hippocampus during placement of the other tetrodes and to later obtain simultaneous recordings from a dendritic layer.

### Data acquisition

Data were acquired using a Digital Lynx system and Cheetah recording software (Neuralynx, Bozeman, Montana). The recording setup has been described in detail previously^9^. Briefly, LFPs from one channel within each tetrode were continuously recorded in the 0.1-500 Hz band at a 2,000 Hz sampling rate. Input amplitude ranges were adjusted before each recording session to maximize resolution without signal saturation. Input ranges for LFPs generally fell within +/-2,000 to +/-3,000 μV. To detect unit activity, signals from each channel in a tetrode were bandpass filtered from 600 to 6,000 Hz. Spikes were detected when the signal on one or more of the channels exceeded a threshold set at 55 μV. Detected events were acquired with a 32,000 Hz sampling rate for 1 ms. Signals were recorded differentially against a dedicated reference channel (see “Surgery and tetrode positioning” section above). Video was recorded through the Neuralynx system with a resolution of 720 x 480 pixels and a frame rate of 29.97 frames per second. Animal position was tracked via an array of red and blue light-emitting diodes (LEDs) on two of the three HS-27-Mini-LED headstages (Neuralynx, Bozeman, Montana) attached to the recording drive.

### Behavioral task and performance

Rats were trained to run laps uni-directionally on a circular track (diameter of 100 cm, height of 50 cm, and width of 11 cm) beginning and ending at a wooden rest box (height of 31 cm, length of 28 cm, and width of 21 cm) attached to the track. 19 small white door bumpers were used to mark potential reward locations on the track. Each door bumper was at a distance of 10 cm (or equivalently 0.199 radians) away from its neighboring door bumpers. For every recording session, there were three stages in the behavioral task: Pre-running, Sample-test, and Post-test. In between task stages, rats rested in the wooden box for 5 minutes. During the pre-running stage, rats ran 4 or 6 laps on the track without receiving any food reward. The Sample-test stage consisted of 8 pairs of sample and test trials. In sample trials, a pseudo-randomly chosen goal location was marked by a yellow plastic cue placed on top of the corresponding door bumper. Of the 19 door bumpers that were on the track, only the 10 door bumpers that were the furthest distance from the wooden box were included as potential reward locations. After a sample trial, rats would rest for 30 seconds in the wooden box, during which time all cues would be removed by the experimenter (i.e., the yellow plastic cue was removed from the goal location, and potential olfactory cues were removed). Then, rats would run a test trial during which they were required to run toward the unmarked reward location and stop there. One of two tones (2 and 6 kHz sine wave with a duration of 0.25 s) signaling a correct choice or an error would sound if rats stopped at the correct reward location or at a different location. The two tones were randomly assigned to signal a correct or error choice for each rat. Rats received a food reward (i.e., Froot Loops piece) from the experimenter after the correct tone sounded. Rats had to stop only at the correct goal location to receive a reward. If rats first stopped at a wrong location, no reward would be given in the trial even if they subsequently stopped at the correct goal location. Paired sample-test trials were separated by an approximately 1-minute inter-trial interval and were repeated 8 times. During the following Post-test phase, rats ran 4 or 6 more test trials to check retention of their memory for the session’s goal location. The goal location remained constant throughout a recording session but was not repeated across successive recording sessions. Rats typically performed one session per day, but 3 out of the 4 rats performed two sessions per day (one in the morning and one in the afternoon) on days when their cell yields were particularly high. A dynamic state-space model was implemented using WinBUGS algorithms^50, 51^ to estimate rats’ probability of making a correct response during Test and Post-test trials and to compute confidence intervals for the estimated probability. Baseline probability in the model was set to be 0.2 since rats chose one of the 5 door bumpers closest to a goal location in most recording sessions.

### Histology

Histological sections were prepared in the following manner to verify tetrode positions at the end of experiments. Rats were given a lethal dose of Euthanasia III solution via intraperitoneal injection. Rats then received intra-cardiac perfusion with phosphate-buffered saline followed by formalin to fix brain tissue. Rat brains were then sliced into 30 µm coronal sections and stained with cresyl-violet to confirm final tetrode positions in CA1. In one rat, a poor perfusion precluded localization of each individual tetrode, but damage from the drive implantation was observed in the cortex overlying CA1, and tetrode depths and signals were consistent with placement in CA1.

### Spike sorting and unit selection

Detected spikes were manually sorted using graphical cluster-cutting software (MClust; A.D. Redish, University of Minnesota, Minneapolis). Spikes were sorted using two-dimensional projections of three different features of spike waveforms (i.e., energies, peaks, and peak-to-valley differences) from four channels. Units with mean firing rates of more than 5 Hz were identified as putative fast-spiking interneurons and were excluded from further analysis. To calculate position tuning for the remaining units, numbers of spikes were divided by time of occupancy at each 0.07-radian position bin on the circular track, followed by smoothing with a Gaussian kernel (standard deviation of 0.14 radians). The running speed for each position (X_n_) was estimated by taking the difference between the preceding position (X_n-1_) and the following position (X_n+1_) and dividing by the elapsed time (2*1/position sampling frequency). Only times when rats moved at speeds above 5 cm/s were included to estimate position tuning. For all single units, position tunings during and after pre-running were compared (Supplementary Fig. 1a-d). Consistent with previous studies^52, 53^, the proportion of units active at a goal location increased after learning. Since over-representation of a goal location could potentially bias our decoding analysis (see “Bayesian decoding analysis” section below), units with peak firing rates above 0.5 Hz at a goal location after pre-running were randomly downsampled to match the number of units with peak firing rates above 0.5 Hz elsewhere on the track, excluding the positions that corresponded to the rest box.

### Bayesian decoding analysis

To translate ensemble spiking activity into angular positions on the circular track, a Bayesian decoding algorithm was implemented^54^. Decoding was performed using a 40-ms sliding time window that shifted 10 ms at each step. The probability of a rat being at position x, given the number of spikes n from each unit recorded in a time window, was estimated by Bayes’ rule:

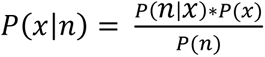

where P(n|x) was approximated from the position tuning of each unit after pre-running (i.e., during sample-test and post-test stages). The approximation assumed that the position tuning of individual units were statistically independent and the number of spikes from each unit followed a Poisson distribution^54^. Prior knowledge of position, P(x), was set to 1 to avoid decoding bias to any particular location on the track. The normalizing constant, P(n), was set to ensure P(x|n), or posterior probability, summed to 1. In order to validate the decoding results, the decoder trained from position tunings after pre-running was used to decode ensemble spiking activity during pre-running. Decoding error was defined as the distance between the decoded position with maximum-likelihood and the rat’s true position. Cumulative decoding errors for each recording session and their average confusion matrices confirmed decoding accuracy of the Bayesian decoder (Supplementary Fig. 1e,f).

### Detection of place cell sequences

Place cell sequences were characterized by continuous decoding of ensemble spiking activity that met the following criteria: a sequence included at least 6 consecutive time bins (90 ms) that contained a spike, estimated positions between adjacent time bins did not exceed 1.4 radians, distance between the first and last estimated position within a sequence was more than or equal to 0.07 radians, and there were at least 3 different cells and 5 spikes within a sequence. To compute the slope for place cell sequences, a circular-linear regression line was fitted to the posterior probability distribution^55^. For a sequence to be included in further analyses, at least 60% of the total posterior probability needed to be no more than 0.35 radians away from the fitted trajectory line. In addition, minimal distance between the fitted trajectory and rats’ true position had to be less than or equal to 0.35 radians. Sequences were analyzed as rats approached their stop locations. Sequences with positive (Fig. 3 and 4, n = 2838 and 2850, respectively) and negative (Supplementary Fig. 4, n = 1611 for sample phase and n = 1558 for test phase) slopes were analyzed separately. Sequences that began before the stop location but contained decoded locations that extended beyond the stop location were also included. Sequences were not analyzed as animals ran on the track during the “Pre-running” stage because animals had been trained to not stop along the track during this stage (i.e., just to continue running past all potential reward sites).

For almost all analyses throughout the paper, place cell sequences were detected based only on the sequential structure in spike trains, as described above. For the analysis in which sequences were detected within individual theta cycles, the following methods were applied before sequential structure was detected using the same criteria as described above. Individual theta cycles were cut at the theta phase with the lowest number of spikes (typically at the peak or close to the peak) from all recorded CA1 cells during that session (as in our previous study^9^). Bayesian decoding was performed for those theta cycles containing spikes from at least 3 different cells and at least 5 spikes within the theta cycle. Also, the rat’s mean running speed during the time of the theta cycle was required to exceed 5 cm/s (≈0.1 radians/s).

### LFP signal processing

Theta (6-10 Hz), slow gamma (25-55 Hz), and fast gamma (60-100 Hz) powers were estimated from the squared amplitude of the Hilbert transform of their corresponding bandpass-filtered signals. Next, power was normalized across the time when rats were moving (>= 5 cm/s) outside of the rest box, and the normalization was done separately for each frequency band and recording session. Among simultaneously recorded CA1 tetrodes with place cells, the tetrode with the highest raw power in the theta band was selected for normalized power (Supplementary Fig. 11) and phase (Fig. 5 and 8) estimates. Theta, slow gamma, and fast gamma phases were estimated using the angles of their Hilbert transformed signals.

### Categorization of place cell locations

Positions from the beginning of the track to the reward location were divided into 3 equally spaced segments (i.e., Early, Middle, and Late). Within each place cell sequence, active place cells with peak firing positions inside each position segment were sorted into corresponding categories (Fig. 5). This analysis was designed to test whether the reported difference in slopes between error and correct trials (i.e., Fig. 3 and 4) was explained by sequences during error trials starting at later theta phases or terminating at earlier theta phases than during correct trials. Thus, only spikes from place cell sequences that were active during the period from 10 to 5 positions before the goal location (i.e., the approximate period showing significant differences in slopes between correct and error trials; see Fig. 3e and 4e), were used to obtain the theta phase estimates shown in Fig. 5.

### Detection of SWRs and replay events

Using LFPs recorded from CA1 tetrodes with place cells, SWRs were detected when rats were in the rest box. The detection method followed that described in a previous study^56^. In brief, LFPs were first bandpass filtered between 150 and 250 Hz, followed by a Hilbert transform to estimate the instantaneous wave amplitude of the filtered LFPs. After smoothing the amplitude with a Gaussian kernel (standard deviation of 25 ms), a SWR event was detected if the amplitude exceeded 5 standard deviations above the mean for at least 50 ms. Boundaries of detected SWR events were adjusted to first crossings of the mean amplitude. Next, the Bayesian decoder described previously (see “Bayesian decoding analysis” section) was applied to ensemble spiking activity within SWR events. To evaluate how well a posterior probability distribution of positions across time resembled a behavioral trajectory on the circular track, the same circular-linear regression method as described above was used to fit a line to the probability distribution (see “Detection of place cell sequences” section). The corresponding r^2^ values were used to quantify replay quality following previously published procedures^12, 48^. A SWR event was classified as a forward or reverse event if the slope of the fitted line was positive or negative, respectively (Supplementary Fig. 5). A replay event was identified if a SWR event’s r^2^ value was above 0.5, and there were fewer than 20% of time bins without any spikes. Boundaries of a replay event were adjusted to the first and last time bins with spikes inside a SWR event. The adjusted duration of a replay event was required to exceed 50 ms to be included for further analysis. Termination bias of replay events at a goal location was evaluated by aligning the last several time bins of each replay event and different goal locations from each session using a method adapted from a previous study^18^ (Fig. 7, Supplementary Fig. 6, and Supplementary Fig. 7). A shuffling method was employed to determine whether termination bias at the goal location was significant. For each replay event, its entire posterior probability across time was circularly shifted 5,000 times by a random distance ranging from 0 to 2π. Summation of the shifted posterior probability across time provided a null distribution to compare against the real data (Fig. 7b,d; Supplementary Fig. 6b,d; and Supplementary Fig. 7b,d). Initiation bias of replay events at a goal location was evaluated using the same method except that the first several time bins of each replay event were aligned instead of the last several time bins (Supplementary Fig. 8).

Replay events were characterized separately for trials 1-4 and for trials 5-8. Across animals and recording sessions, behavioral performance stabilized at an above chance performance level at trial 5 (Fig. 1b). Therefore, rest periods from trials 1-4 and trials 5-8 were assumed to represent rest periods when learning was incomplete and relatively complete, respectively.

### Estimation of gamma phase shifts across successive gamma cycles

To measure gamma phase modulation of spike times across gamma cycles, we estimated gamma phases and theta phases of spike times for place cells that spiked within a detected sequence (based on an analysis used by Zheng and colleagues^9^). The gamma cycle with maximal spiking of all simultaneously recorded cells was defined as cycle 0 (Fig. 8a,b). Cycles occurring before cycle 0 were numbered with decreasing integer values, and cycles occurring after cycle 0 were numbered with increasing integer values. Incomplete cycles at the beginning or end of a sequence were excluded from analyses. The number of cycles analyzed within each sequence was limited to 3 (cycles −1 to 1) for slow gamma and 5 (cycles −2 to 2) for fast gamma as in a previous study^9^. 2-D histograms of gamma phases and theta phases for spikes from each cycle number were plotted using 30° bins, and were smoothed with a box filter (5 × 5 bins). To determine slow gamma phase shifts across successive slow gamma cycles (Figure 8c), circular-linear correlation coefficients between slow gamma phases and cycle numbers were computed using the ‘circ_corrcl’ function from the CircStat Toolbox^57^. The same method was also used for fast gamma phases (Supplementary Fig. 10b). Phase-locking of spike times to fast gamma was measured by the resultant vector length of fast gamma phases of spikes (Figure 8d), pooled across fast gamma cycles for each trial type. The same method was also used to assess slow gamma phase-locking (Supplementary Fig. 10a).

### Data analysis and statistics

Data analyses were performed using custom MATLAB scripts (The Math Works). Paired t-tests, ANOVAs, and repeated measures ANOVAs were performed using standard built-in MATLAB functions (i.e., ‘ttest’, ‘fitlm’ and ‘fitrm’, respectively). A non-parametric circular statistical test was performed to test for differences in the theta phase distributions shown in Fig. 5 using the ‘circ_cmtest’ function from the CircStat Toolbox^57^. Permutation tests (used for the data shown in Fig. 7b,d and Fig. 8c,d) shuffled independent variables (i.e. angular position in Fig. 7b,d and trial type in Fig. 8c,d) 5,000 times to generate a null distribution. A result was considered significant if the observed data exceeded the 97.5th percentile or fell below the 2.5th percentile of the null distribution (i.e., a two-tailed test). All confidence intervals were estimated using a standard built-in MATLAB function ‘bootci’ (n = 5,000 resamples). The experimental design of this study did not involve any subject grouping, and thus data collection and analyses were not performed blind.

### Code availability

The custom MATLAB scripts used in this study are available from the corresponding authors upon reasonable request.

### Data availability

The datasets used in this study are available from the corresponding authors upon reasonable request.

## SUPPLEMENTARY FIGURE LEGENDS

**Supplementary Figure 1.**
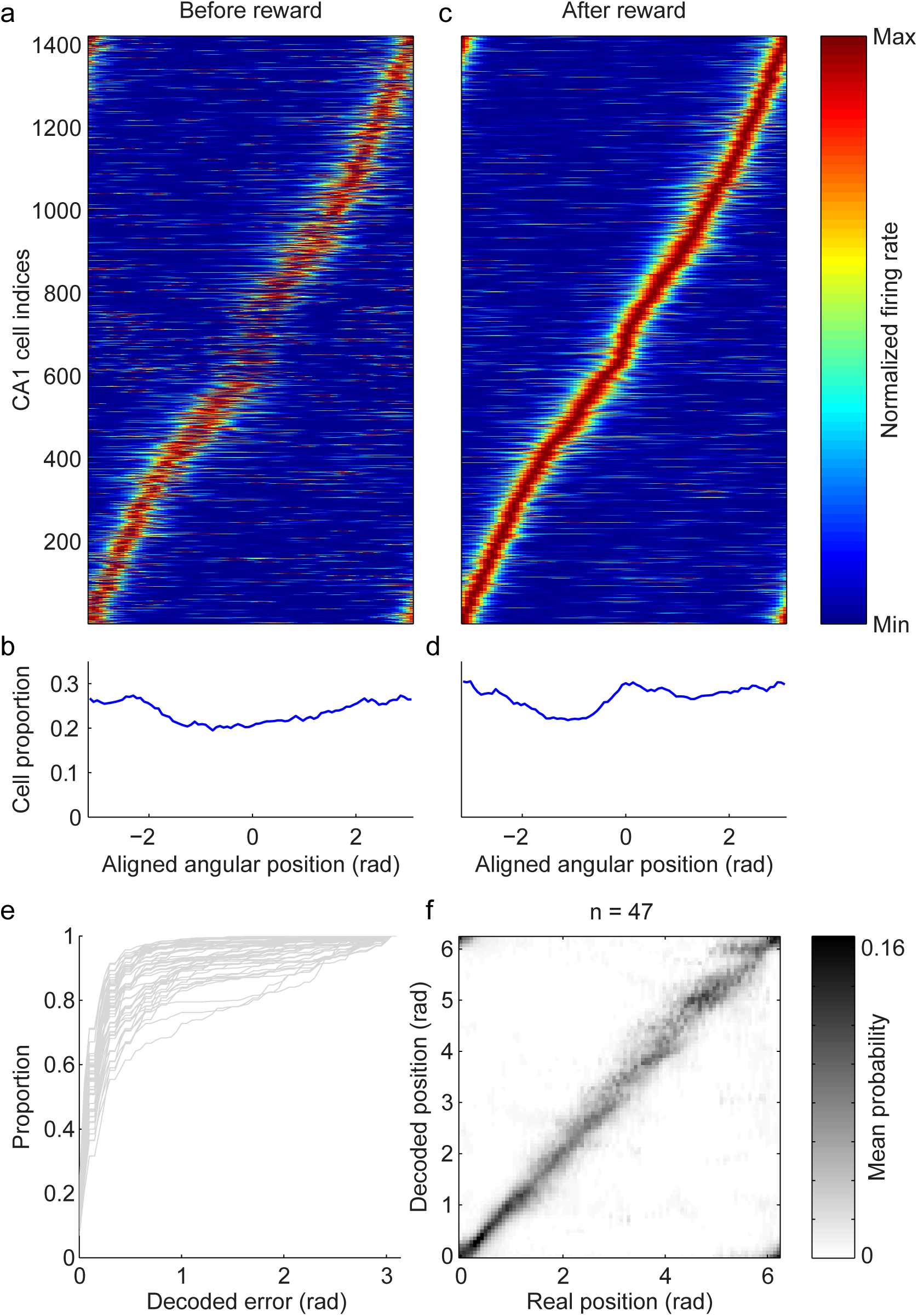
Over-representation of reward location after learning and accuracy of decoding. **a**, Normalized firing rates of CA1 place cells recorded from all sessions are shown for pre-running trials. Place cells were sorted by their peak firing positions after pre-running trials (i.e., during sample, test, and post-test trials). Position was aligned to the reward location on the horizontal axis (i.e., position 0). **b**, Proportion of total cells that exceeded a firing rate threshold of 0.5 Hz across aligned positions in pre-running trials. **c-d**, Same as **a-b** but for sample, test, and post-test trials. **e**, A cumulative graph of decoding errors for each recording session. Each gray trace indicates an individual recording session. **f**, A mean confusion matrix averaged across all recording sessions.

**Supplementary Figure 2.**
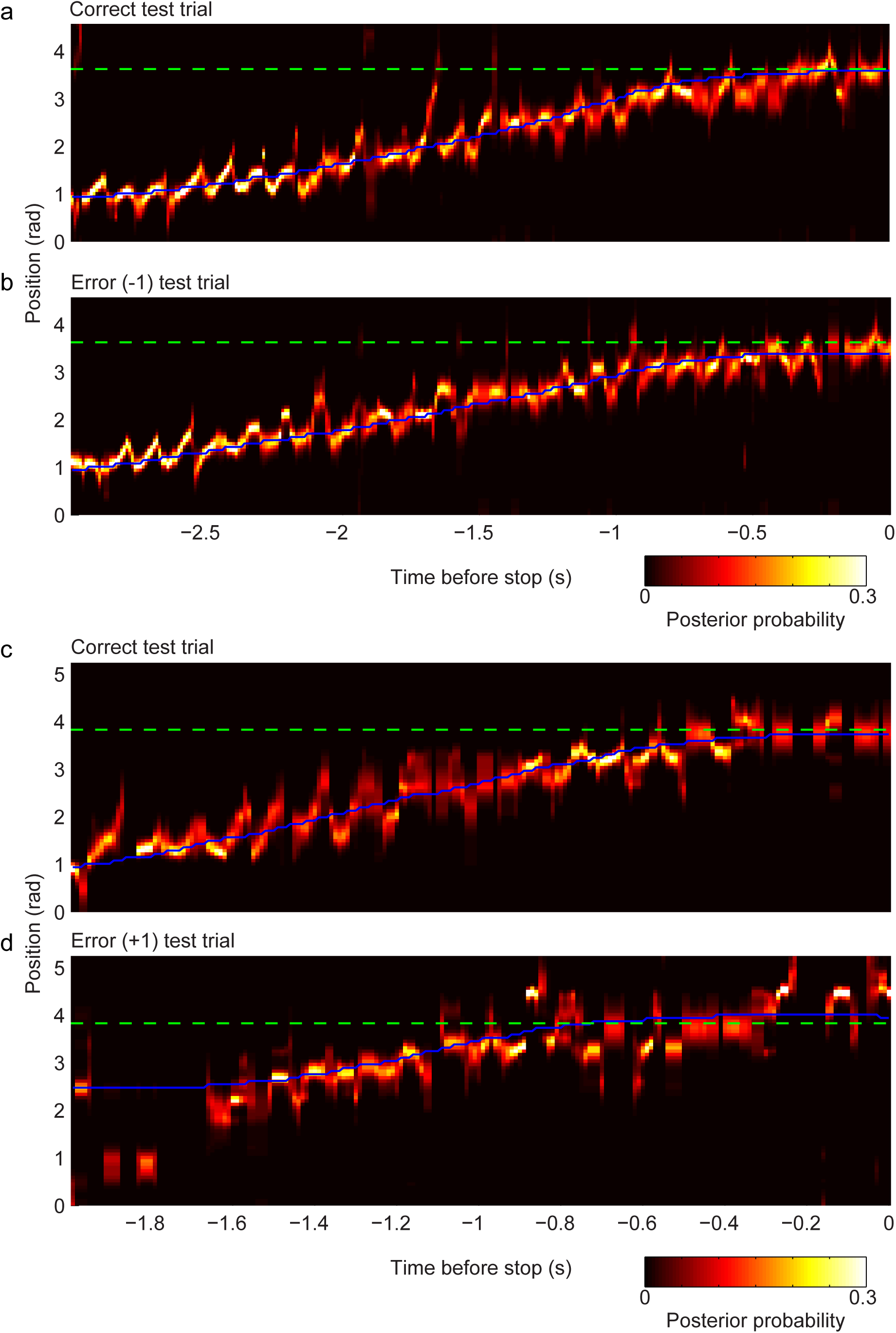
Examples of posterior probability distributions from test trials with different types of errors. **a-b**, Same as Fig. 3 **a-b** but showing a different trial pair example from a different recording session. In this example, the rat stopped one location before the correct reward location in the error test trial. **c-d**, Same as **a-b** but showing a different trial pair example from another recording session in a different rat. In this example, the rat stopped one location later than the correct goal location in the error test trial.

**Supplementary Figure 3.**
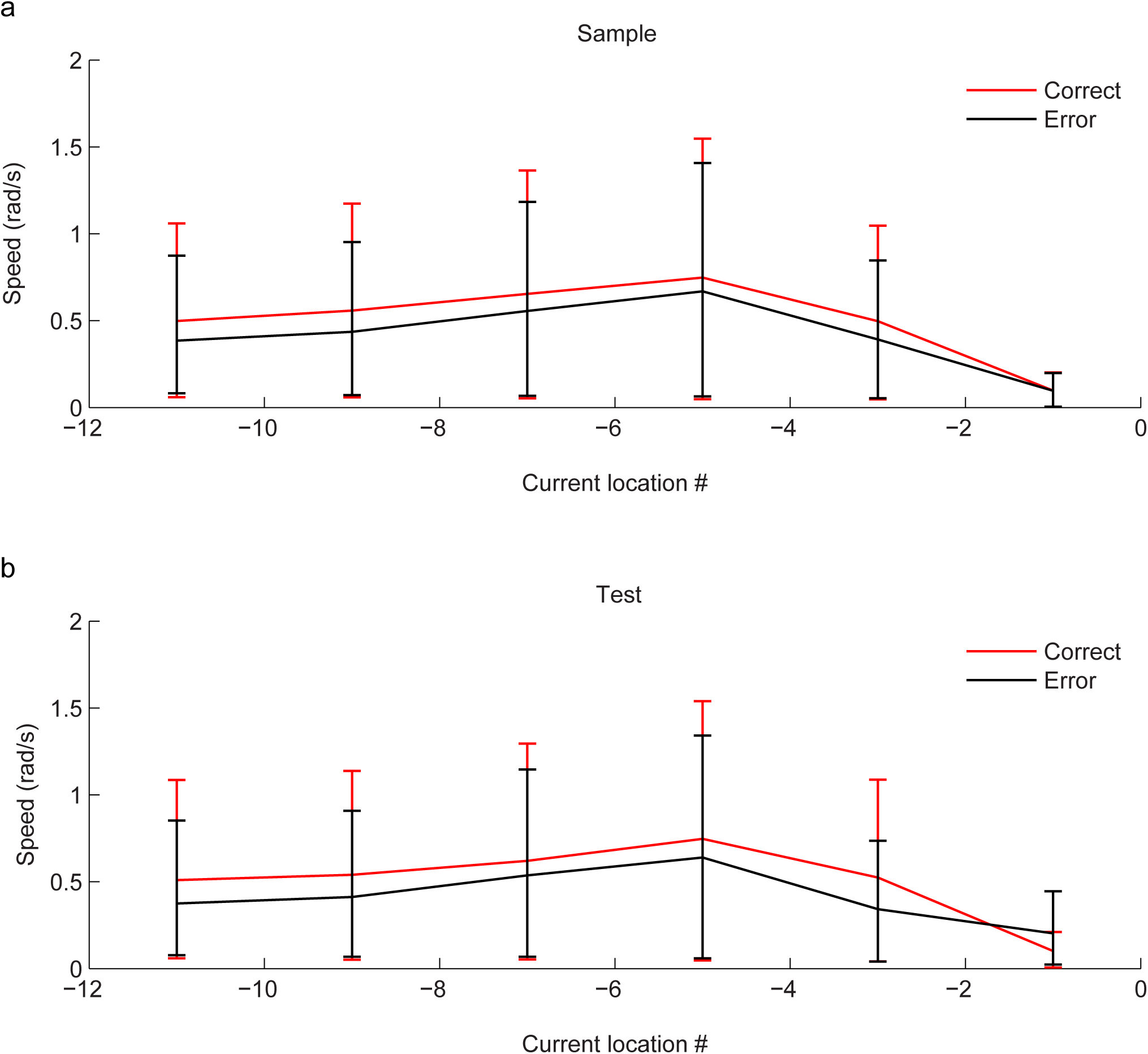
Running speeds during the approach to the stop location. **a**, Running speed estimates during the sample phase of the task are shown across location numbers as rats approached their stop location (indicated as location 0; the stop location was the same as the correct goal location for correct trials and was the incorrect stop location for error trials). Correct trials are shown in red and error trials are shown in black. **b,** Same as **a** but for the test phase.

**Supplementary Figure 4.**
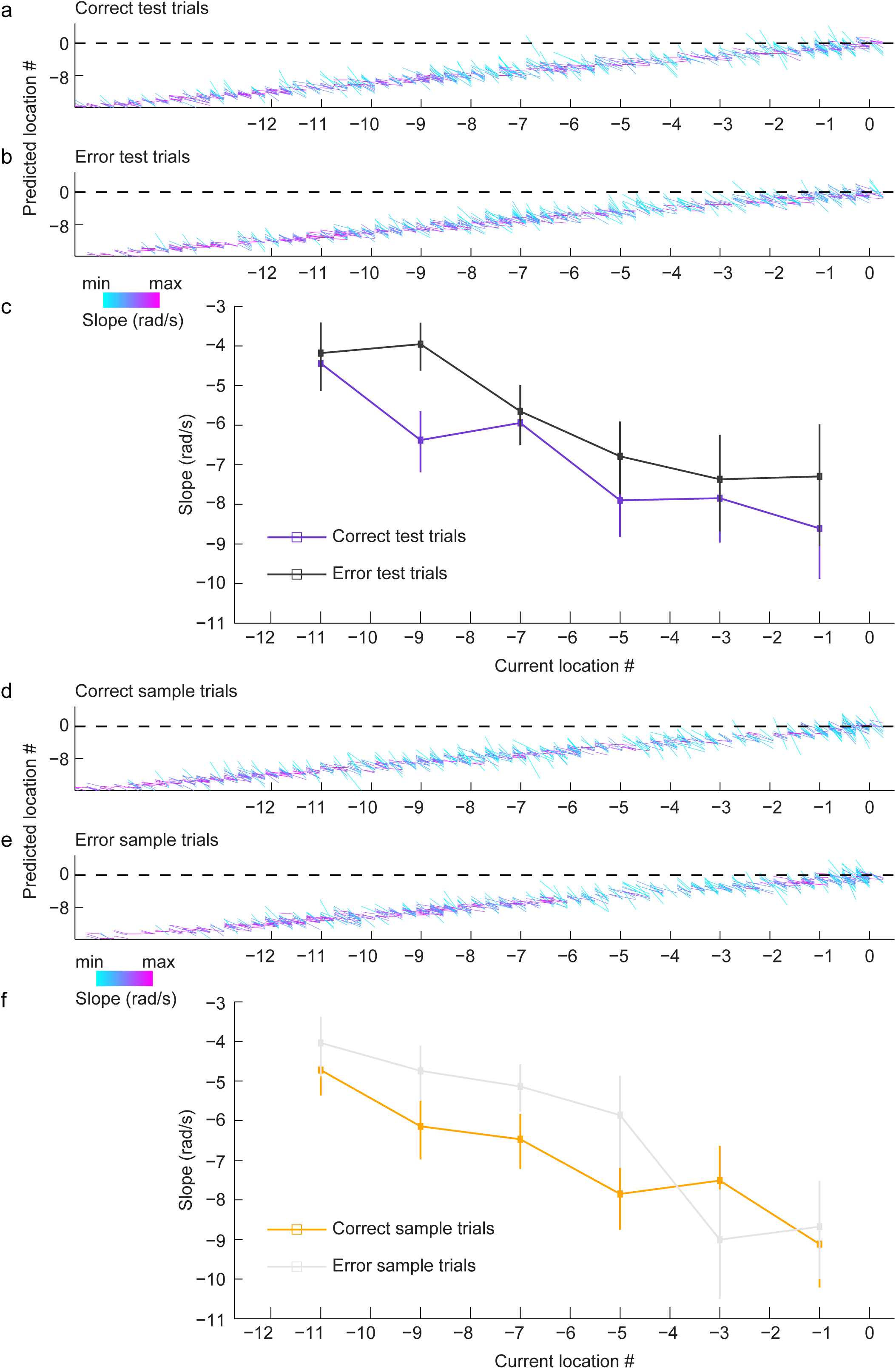
Place cell sequences with negative slopes detected during correct and error trials. **a-c**, Same as Fig. 3 **c-e** but for sequences with negative slopes. **d-f**, Same as **a-c** but for negative slope sequences from the sample phase.

**Supplementary Figure 5.**
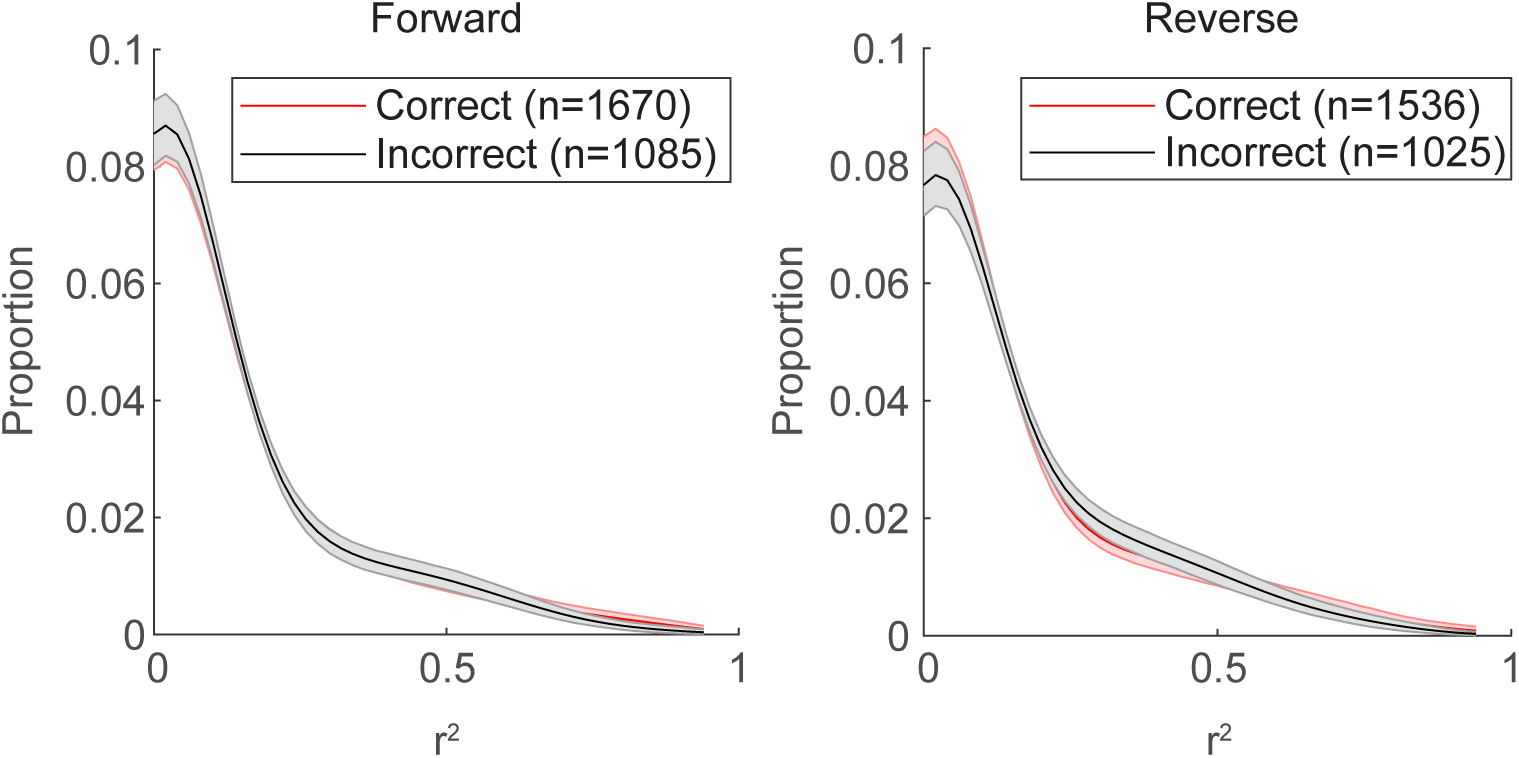
Forward and reverse replay fidelity was similar between rest periods from correct and error trials. Distributions of r^2^ values for forward (left panel) and reverse (right panel) replay events for all detected SWR events in correct (red) and error (black) trials. Shaded error bars indicate 95% bootstrapped confidence intervals.

**Supplementary Figure 6.**
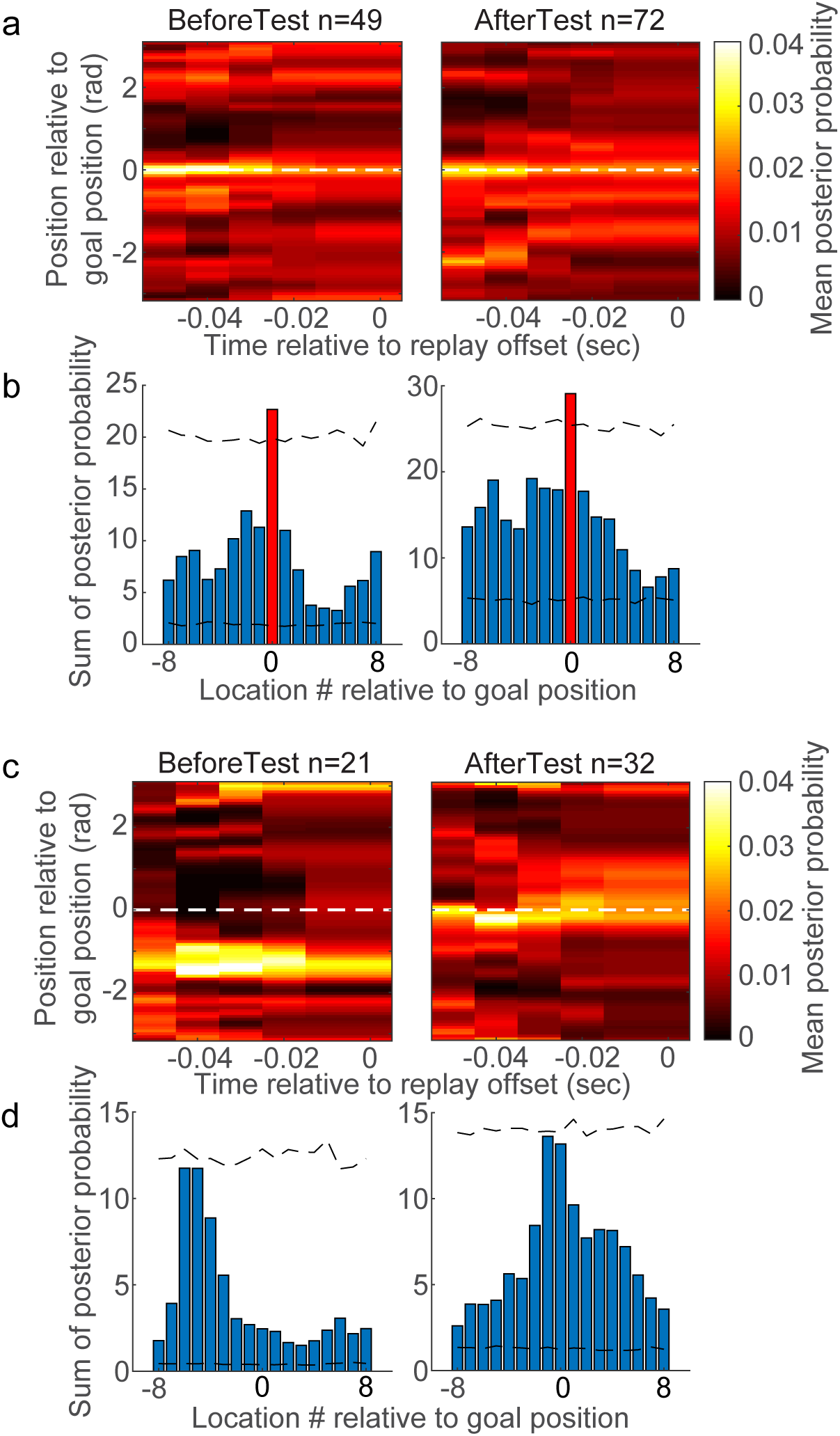
A bias for replay events to terminate at the correct reward location occurs in rest periods before and after the test phase during correct trials but not error trials. **a**, Mean posterior probabilities of replay events in rest periods of correct trials before the test phase (left panel) and after the test phase (right panel). Replay events were aligned to the correct goal location on the y-axis (i.e., location 0) and the time of replay event offset (time = 0) on the x-axis. **b**, The sum of posterior probability across time for each location number relative to the correct goal location. Dashed black lines mark 95% confidence intervals of a null distribution generated by randomly shifting positions of each replay event, as in Fig. 7b and d. Red bars mark the location numbers with a posterior probability sum that exceeded the corresponding 95% confidence interval (i.e., location 0, which corresponds to the correct goal location). **c**, Same as **a** but for error trials. **d,** Same as **b** but for error trials. Note that no bars are red because none of the locations’ summed posterior probabilities exceeded the 95% confidence intervals for error trials.

**Supplementary Figure 7.**
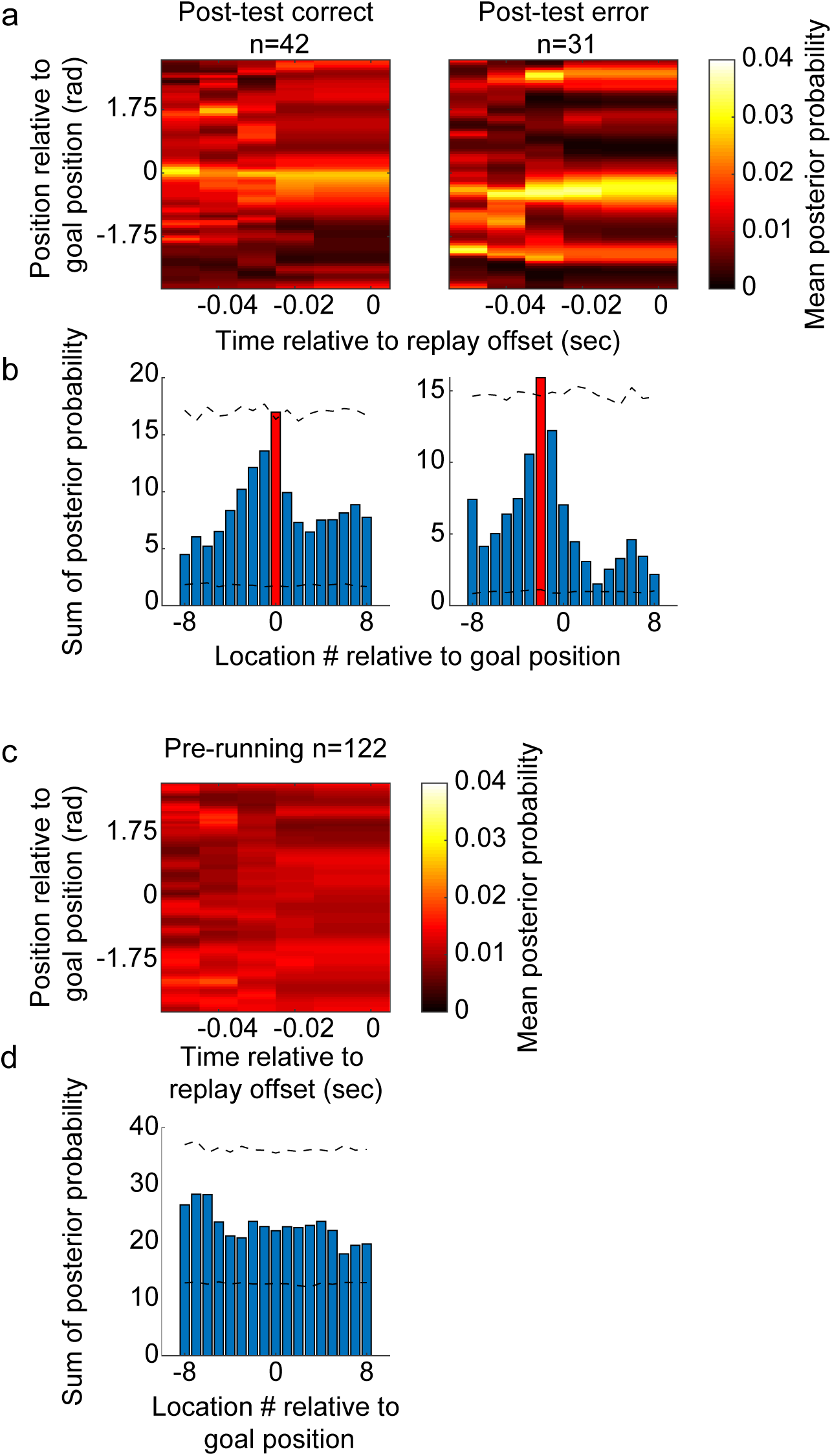
A bias for replay events to terminate at the correct reward location persisted in correct post-test trials but was absent in pre-running trials. **a**, Mean posterior probabilities of replay events in post-test trials. Replay events were aligned to the correct goal location on the vertical axis (i.e., location 0) and the time of replay event offset on the horizontal axis. Correct and error post-test trials are shown in the left and right panels, respectively. **b**, The sum of posterior probability across time for each location number relative to the goal. Dashed black lines mark 95% confidence intervals of a null distribution generated by randomly shifting positions of each replay event, as in Fig. 7b and d. A red bar marks the location number with a posterior probability sum that exceeded the corresponding 95% confidence interval (i.e., location 0, which corresponds to the correct goal location, for correct post-test trials and location −2, which corresponds to an incorrect location, for error post-test trials). **c-d**, Same as **a-b** but for pre-running trials. Since there was no memory test or reward in pre-running trials, all trials were of the same type (i.e., no correct and error trials).

**Supplementary Fig. 8.**
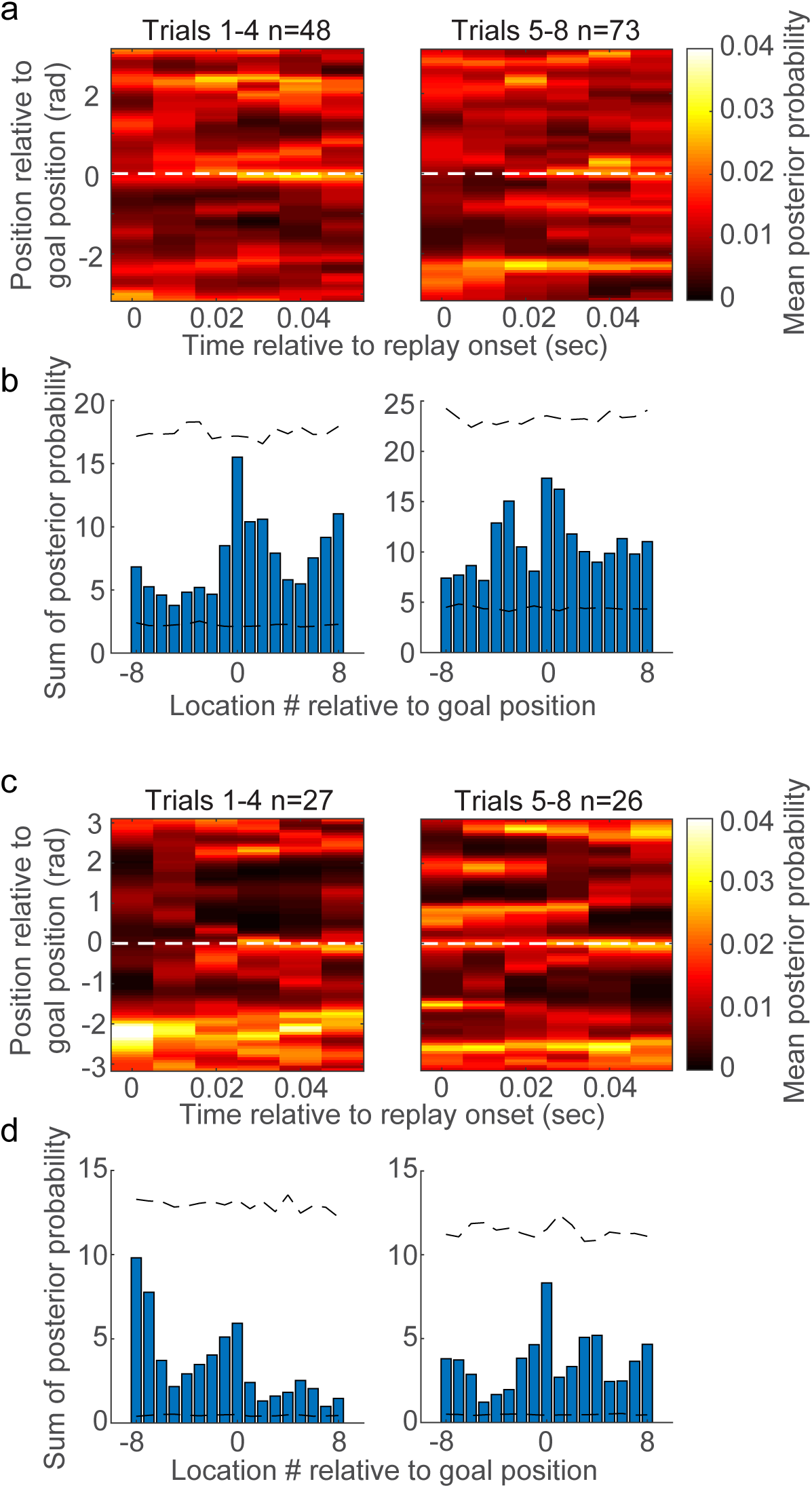
No bias for replay events to start at the correct goal location was observed. **a**, Mean posterior probabilities of replay events in correct trials 1-4 (left panel) and correct trials 5-8 (right panel). Replay events were aligned to the correct goal location on the y-axis (i.e., location 0) and the time of replay event onset on the x-axis. **b**, The sum of posterior probability across time is shown for each location number relative to the correct goal location (i.e., location 0). Dashed black lines mark 95% confidence intervals of a null distribution generated by randomly shifting positions of each replay event (see Methods). None of the locations’ summed posterior probabilities exceeded the 95% confidence intervals. **c-d**, Same as **a-b** but for error trials.

**Supplementary Figure 9.**
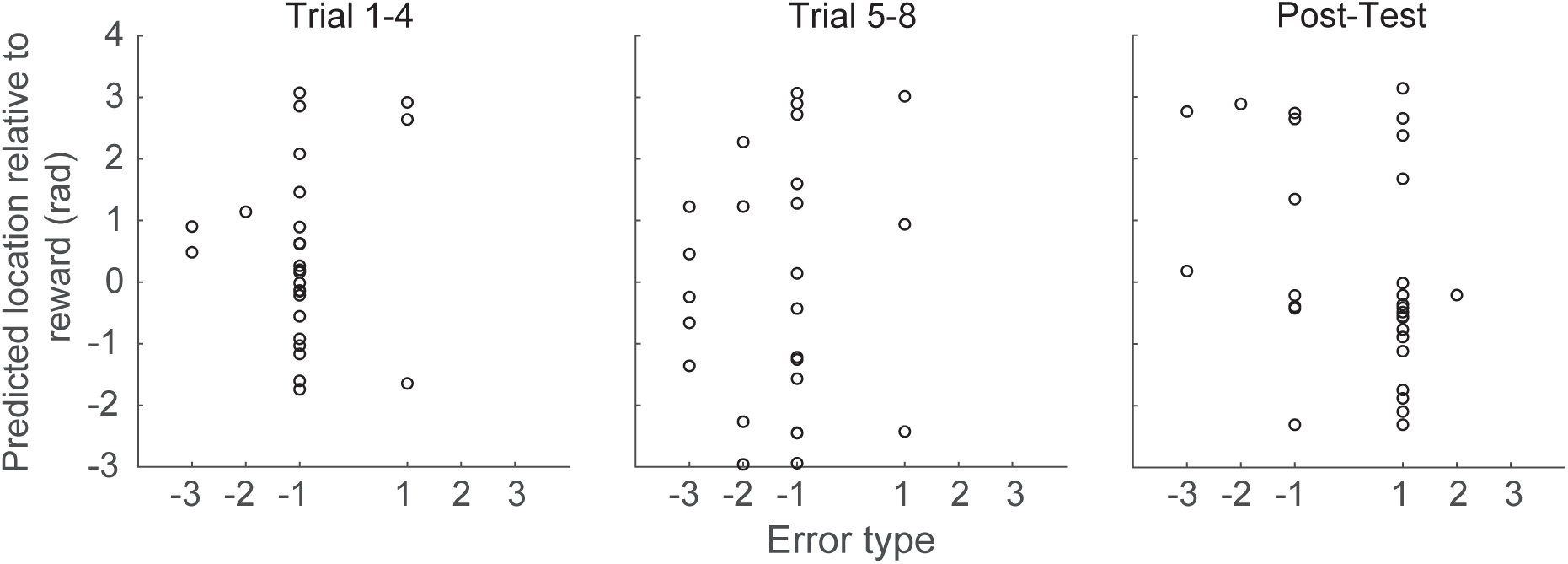
No relationship between locations represented at the end of replay sequences and animals’ actual stop locations was observed during error trials. The y-axis shows the location represented at the end of each replay sequence detected during rest periods of error trials for different error types shown on the x-axis. Error types −3 to −1 indicate that rats stopped at incorrect locations that were 3 to 1 locations before the correct location. Error types 1 to 3 indicate stop locations that were 1 to 3 locations after the correct reward location. Results from errors in trials 1-4, trials 5-8, and post-test trials are shown in left, middle, and right panels, respectively.

**Supplementary Figure 10.**
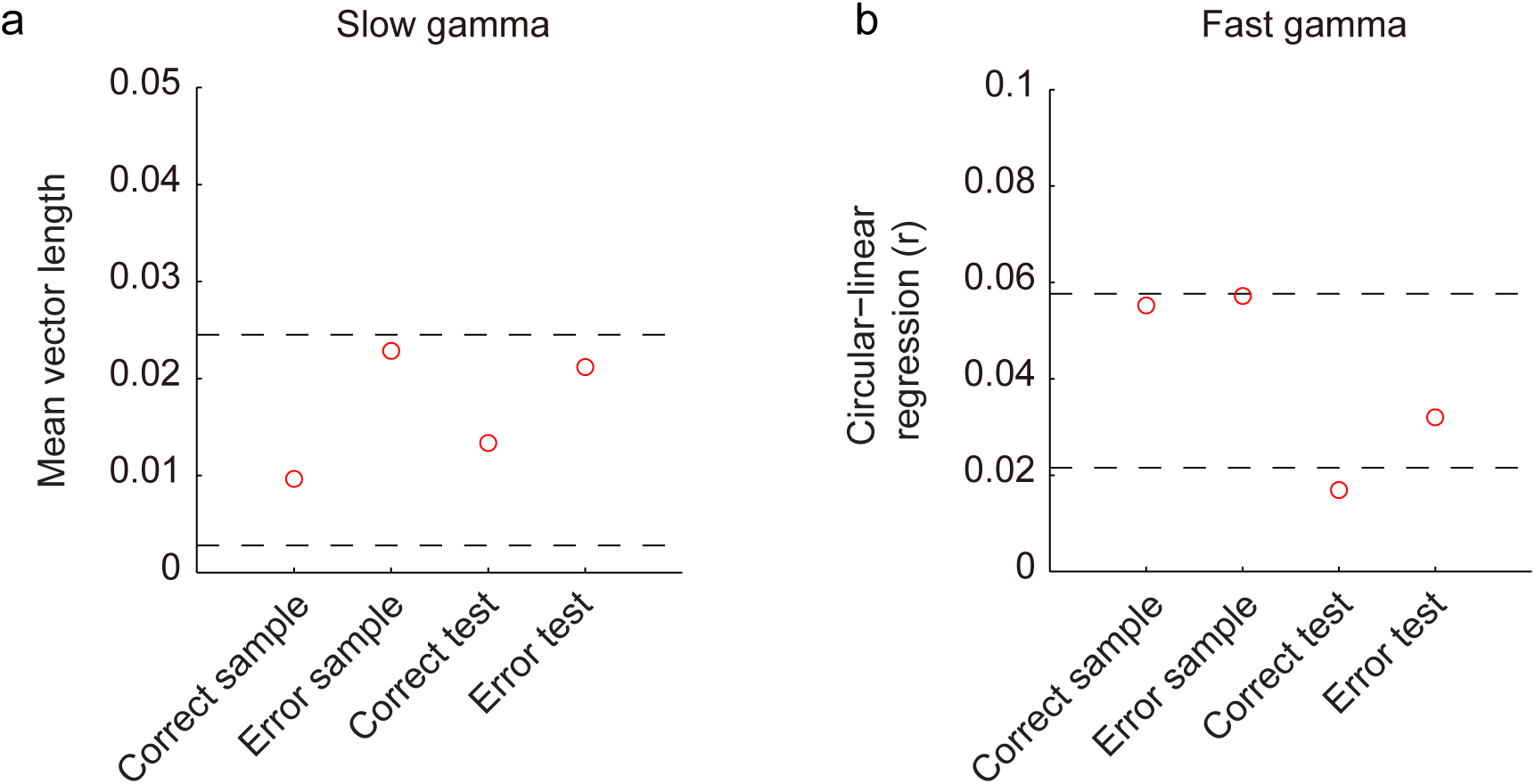
Additional gamma phase analyses. Black dashed lines mark 95% confidence intervals of a null distribution generated by shuffling trial types. **a,** Mean vector lengths of slow gamma phases of place cell spikes for each trial type, pooled across all slow gamma cycles. **b,** Circular-linear regression between fast gamma phases of spikes and cycle number for each trial type. The correct test trials’ value fell outside the confidence intervals, but it is unclear to us how to interpret this finding. These analyses were not planned to test hypotheses suggested by our earlier work^9^ but were requested during review.

**Supplementary Figure 11.**
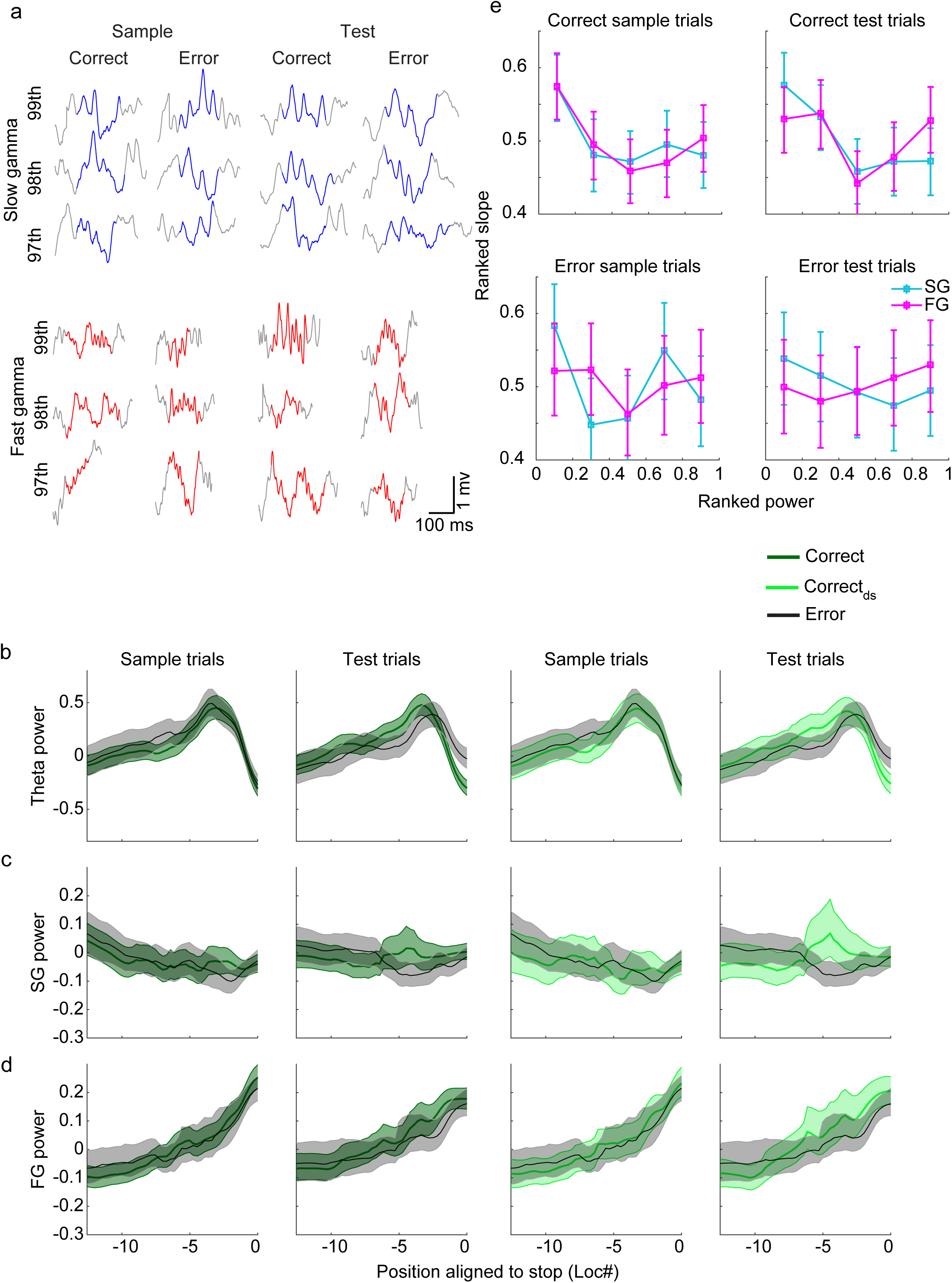
No difference in gamma power was observed between trial types as rats approached a stop location. **a,** Example of detected slow and fast gamma events during sample and test phases of correct and error trials. Shown are examples of raw LFP recordings with power in the 97^th^, 98^th^, and 99^th^ percentiles for each gamma type. **b**, Mean normalized power averaged across the theta frequency band (6-12 Hz) for sample trials (far left panel), test trials (2^nd^ panel), downsampled sample trials (3^rd^ panel), and downsampled test trials (far right panel). Color labels indicate correct, downsampled correct, and error trials (dark green, light green, and black, respectively). Shaded area indicates 95% bootstrapped confidence intervals. **c-d**, Same as **b** but for mean normalized power averaged across the slow gamma (25-55 Hz) and fast gamma (60-100 Hz) frequency bands, respectively. **e**, Relationship between place cell sequence slopes and gamma power for each trial type. Slopes and gamma power estimates for each place cell sequence were ranked and normalized for each trial type. Slow (blue) and fast (magenta) gamma power were plotted separately against sequence slopes. Error bars indicate 95% bootstrapped confidence intervals.

